# Scalable, multimodal profiling of chromatin accessibility and protein levels in single cells

**DOI:** 10.1101/2020.09.08.286914

**Authors:** Eleni P. Mimitou, Caleb A. Lareau, Kelvin Y. Chen, Andre L. Zorzetto-Fernandes, Yusuke Takeshima, Wendy Luo, Tse-Shun Huang, Bertrand Yeung, Pratiksha I. Thakore, James Badger Wing, Kristopher L. Nazor, Shimon Sakaguchi, Leif S. Ludwig, Vijay G. Sankaran, Aviv Regev, Peter Smibert

**Author notes:** Contributed equally (Eleni P. Mimitou, Caleb A. Lareau, Kelvin Y. Chen). Senior author (Shimon Sakaguchi, Leif S. Ludwig, Vijay G. Sankaran, Aviv Regev, Peter Smibert).

## Abstract

Recent technological advances have enabled massively parallel chromatin profiling with <u>s</u>ingle-<u>c</u>ell <u>A</u>ssay for <u>T</u>ransposase <u>A</u>ccessible <u>C</u>hromatin by <u>seq</u>uencing (scATAC-seq) in thousands of individual cells. Here, we extend these approaches and present <u>A</u>TAC with <u>S</u>elect <u>A</u>ntigen <u>P</u>rofiling by <u>seq</u>uencing, ASAP-seq, a tool to simultaneously profile accessible chromatin and protein levels in thousands of single cells. Our approach pairs sparse scATAC-seq data with robust detection of hundreds of cell surface and intracellular protein markers and optional capture of mitochondrial DNA (mtDNA) for clonal tracking, thus concomitantly capturing three distinct modalities in single cells. Importantly, ASAP-seq uses a novel bridging approach that repurposes antibody:oligo conjugates designed for existing technologies that pair protein measurements with single cell RNA-seq. We demonstrate the utility of ASAP-seq by revealing coordinated and distinct changes in chromatin, RNA, and surface proteins during native hematopoietic differentiation, peripheral blood mononuclear cell stimulation, and as a combinatorial decoder and reporter of multiplexed perturbations in primary T cells.

## INTRODUCTION

The recent explosion of technologies allowing detailed phenotypic measurements of single cells in high-throughput has made the dissection of cell types and states in complex tissues accessible to most researchers. While measurement of single modalities has been highly informative for phenotyping, new techniques that allow detection of multiple modalities of information from single cells continue to be developed^1–4^.

Multi-modal approaches couple sparse comprehensive measurements with more robust directed measurements that report on known cell types or states. For example, CITE-seq^5, 6^ and REAP-seq^7^ couple scRNA-seq with detection of surface proteins. In these methods, oligo-labeled antibodies detect highly abundant and well-characterized surface protein markers, which complement the relatively sparse scRNA-seq signal and enable more robust cell type discrimination, relating different levels of gene regulation and connecting to a rich body of work on phenotypes at the protein level. However, while mRNA and protein are the products of gene expression, their detection (or lack thereof) at a snapshot in time do not suffice to decipher the underlying regulatory mechanisms at their respective genomic loci.

The chromatin architecture of a cell is an early phenotypic readout that highlights regulatory mechanisms that control some of the earliest steps in gene expression, in instances allowing detection of the earliest cellular responses to stimuli or developmental decisions, and identification of poised states^8^. In particular, the <u>A</u>ssay for <u>T</u>ransposase-<u>A</u>ccessible <u>C</u>hromatin by <u>seq</u>uencing applied to single cells (scATAC-seq) is a recent but widely-used method to obtain a genome wide snapshot of chromatin accessibility, signatures of active transcription and even transcription factor binding^9, 10^. Several methods have recently been developed for the capture of mRNA together with chromatin accessibility in single cells and help correlate chromatin accessibility with gene expression, as well as layer mRNA expression data on top of sparse ATAC-seq data^8, 11–13^. While having transcript and chromatin accessibility data from the same single cells is valuable, the ultimate step of gene expression, in most cases, is regulation of protein levels, and much of our understanding of cell function is associated with such changes. Moreover, changes to protein levels and modifications can happen in ways that are not coupled to transcription and operate at fast time scales, thus preceding changes to regulatory mechanisms, such as chromatin accessibility. Motivated by the recent demonstration that fixed and permeabilized whole cells yield scATAC profiles of comparable quality to traditional fresh nuclear preparations^14^, we sought to combine protein detection with scATAC-seq.

Here, we report <u>A</u>TAC with <u>S</u>elect <u>A</u>ntigen <u>P</u>rofiling by <u>seq</u>uencing (ASAP-seq), a method that enables robust detection of cell surface and intracellular proteins using oligo-labeled antibodies together with high-throughput scATAC-seq. ASAP-seq takes advantage of existing oligo-labeled antibody reagents used for CITE-seq, Cell Hashing, and related technologies, circumventing the need for additional specialized components. Importantly, unlike co-assays of RNA and chromatin, where there is a trade off between enzymatic steps with vastly different requirements, we leverage an approach (as in CITE-seq^5, 6^) that utilizes the enzymatic steps of the parent assay to detect multiple modalities, to ensure high quality across both. Moreover, ASAP-seq is compatible with recent methods designed to detect mtDNA genotypes for lineage tracing and study of mitochondrial diseases^14, 15^. To demonstrate the utility of ASAP-seq, we applied it to the study of human hematopoiesis, where the combination of single cell chromatin accessibility, hundreds of surface marker profiles, and mtDNA-based lineage tracing allowed us to resolve bone marrow heterogeneity and composition. Separately, in a model of immune cell stimulation, we combined ASAP-seq with CITE-seq to reveal the distinct layers of regulation of protein, mRNA levels, and chromatin accessibility. Finally, in a multiplexed perturbation assay in primary T cells, ASAP-seq disentangles chromatin and protein phenotypes associated with specific signalling pathways.

## RESULTS

### Development and validation of ASAP-seq

To develop ASAP-seq, we built on mtscATAC-seq^14^, a droplet-based scATAC-seq method that jointly profiles chromatin accessibility and mtDNA with high coverage in thousands of single cells. In mtscATAC-seq, the retention of mitochondria in fixed whole cells allows their genomes to be tagmented and subsequently sequenced. We reasoned that the fixation and permeabilization before Tn5 transposition would also result in the retention of cell surface markers, enabling their detection with oligo-conjugated antibodies, as demonstrated with CITE-seq and related technologies (**Fig. 1a**)^5–7^. To test whether surface marker detection is compatible with the mtscATAC-seq workflow, we stained peripheral blood mononuclear cells (PBMCs) with fluorophore-labeled antibodies against CD3 and CD19, and performed flow cytometry to measure fluorophore intensity at subsequent steps of the protocol (**Extended Data Fig. 1a**). While permeabilization of fixed cells had a minimal impact on signal intensity, the additional 1 hr incubation at 37°C to mimic the transposition step led to some loss of intensity. However, the separation to background remained distinct, suggesting the workflow to be compatible with antibody-based protein detection.

**Figure 1.**
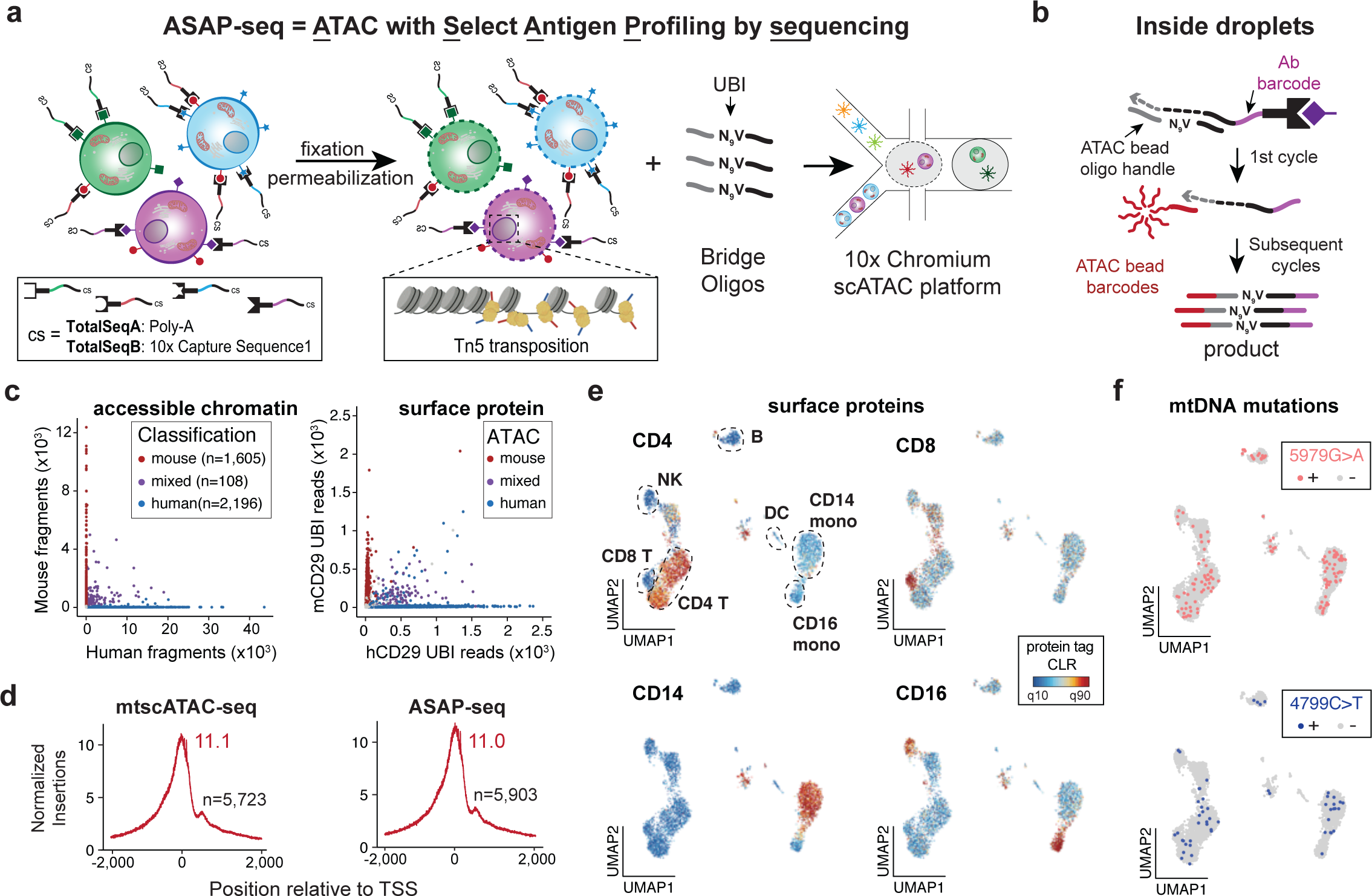
ASAP-seq incorporates protein detection in scATAC-seq workflows. a. Schematic of the cell-processing steps that allow retention and profiling of cell-surface markers jointly with chromatin accessibility. Cells are stained with oligo-conjugated antibodies before fixation, permeabilization and transposition with Tn5. **b.** In droplets, bridge oligos spiked into the barcoding mix promote templated extension of the antibody tags during the first cycle of amplification rendering them complementary to bead-derived barcoding oligos. Extended antibody tags are subsequently barcoded together with the transposed chromatin fragments. **c.** Species mixing experiment using the Pre-SPRI approach; number of unique nuclear fragments (left) and protein-tag counts (right) associated with each cell barcode. Points are colored based on species classification using ATAC-derived fragments (97.4% agreement by assignment; all but 1 discrepancy was an errant doublet versus singlet classification) **d.** TSS enrichment scores of mtscATAC-seq without (left) or with concomitant protein tag capture (right). n indicates the number of cells profiled. **e.** UMAP showing chromatin accessibility-based clustering of PBMCs stained with a 9-antibody panel, with selected markers highlighted. Color bar: protein tag centered log-ratio (CLR) values. **f.** Cellular distribution of two most commonly detected mtDNA mutations in the population. Thresholds for + were 5% heteroplasmy based on empirical density.

We next devised an approach for protein detection that leverages existing validated reagents in broad use. The most parsimonious approach to incorporating protein detection into scATAC-seq would be to design antibody:oligo conjugates with sequences complementary to the oligos that append cell barcodes to tagmented fragments. However, the large existing catalog of commercial antibody:oligo conjugated products designed for scRNA-seq applications (TotalSeq^TM^ products by BioLegend) motivated us to devise a molecular bridging approach, wherein a short oligo added to the reaction mix bridges the interaction between the antibody tag and the barcoding bead oligo in droplets (**Fig. 1b** and **Extended Data Fig. 1b**). The 3’ blocked bridge oligo serves solely as a template for extension of the antibody tag during the initial amplification cycles. The extended product acquires the sequence necessary to anneal to the bead-derived barcoded oligo and is linearly amplified during the subsequent cycles along with accessible chromatin fragments. To allow tag molecule counting when TotalSeq^TM^-A (TSA) products are used, we introduced a Unique Bridging Identifier (UBI) sequence via the <u>B</u>ridge <u>O</u>ligo for TotalSeq^TM^-A (BOA) (**Extended Data Fig. 1b**). As each antibody-derived oligo can only be bridged once, the UBI serves as a proxy for a Unique Molecular Identifier (UMI) and allows individual molecules to be counted. Alternate TotalSeq^TM^ formats (*e.g.*, TotalSeq^TM^-B (TSB) that already contain UMI sequences^16^) do not require UBIs in their bridge oligos (**Extended Data Fig. 2a**).

**Figure 2.**
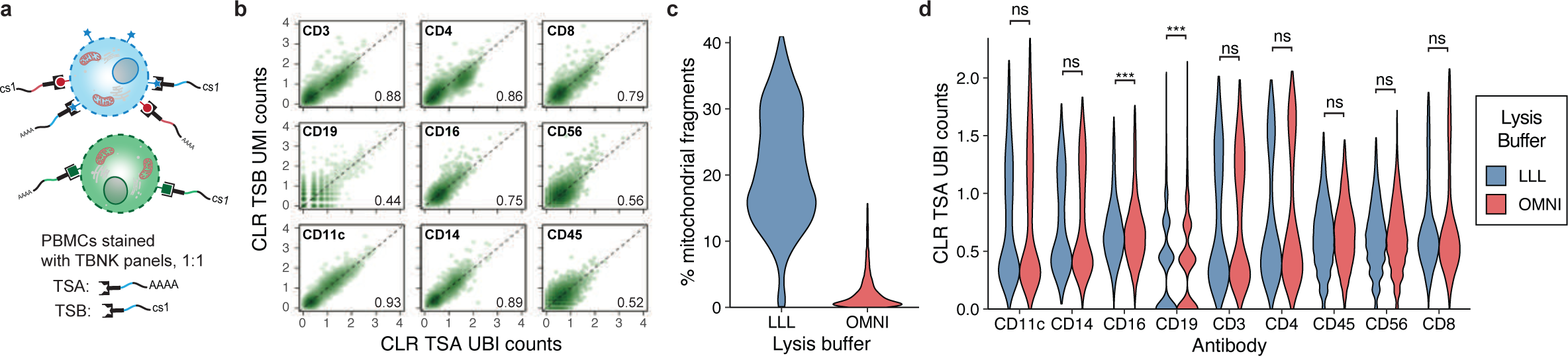
ASAP-seq enables a modular and versatile multi-omics toolkit. a. Schematic of experimental design. PBMCs were stained with TBNK panels of the TSA or TSB format at a 1:1 ratio, followed by fixation and permeabilization under mild (LLL) or strong conditions (OMNI). **b.** Pairwise comparison of centered log-ratio (CLR) normalized TSA and TSB counts for indicated antibodies under mild lysis conditions (n=4,748 cells). Counts were collapsed for unique molecules using UBIs (TSA panel) or UMIs (TSB panel). **c.** Distribution of percent of mtDNA fragments retained in the library under the two lysis conditions. **d.** Comparison of CLR normalized TSA counts for indicated proteins under the two tested lysis conditions. Statistical comparisons are Wilcoxon rank sum test with Bonferroni adjusted p-values (ns = not significant; \**p_adj_* < 0.05; ** *p_adj_* < 0.01; *** *p_adj_* < 0.001).

To benchmark ASAP-seq, we stained a 50:50 mix of human (HEK-293T) and mouse (NIH-3T3) cells with TSA human and mouse-specific anti-CD29 antibodies (**Supplementary Table 1, tab ’mixed species’**), followed by fixation, permeabilization, and transposition prior to barcoding in droplets in the presence of 0.5 µmoles of BOA in the reaction mix (**Methods**). Assigning single cells as mouse, human or multiplet by the number of reads mapping to the respective genomes or by tag identity yielded consistent results (**Methods**), demonstrating the specificity of protein detection in this assay (**Fig. 1c** and **Extended Data Fig. 1c**). In this experiment, two identical barcoding reactions were run in two separate lanes to further test tag library preparation with alternate approaches; ’Pre-SPRI’, where the input for tag indexing is 10% of the purified fragments after emulsion breakage or ’Post-SPRI’, where the input is prepared from the supernatant fraction from the SPRI purification step (**Methods**). In both instances, we observed no substantial changes in ATAC fragment or protein tag complexity, suggesting that either fraction (or a combination of both) can be used to prepare the tag libraries (**Extended Data Fig. 1d,e**). Nevertheless, despite not seeing substantial differences between the two approaches, we opted for the post-SPRI approach for all subsequent assays to retain as many molecules as possible in the fraction used for the generation of ATAC-seq libraries.

To identify differentially expressed proteins and perform additional technical benchmarking and optimization, we applied ASAP-seq to PBMCs stained with a TBNK panel (BioLegend, **Supplementary Table 1, tab ’TBNK’**) that assesses surface expression of 9 major immune cell markers. Concomitantly, we ran a matched unstained sample to assess the impact of antibody tags on scATAC-seq data quality. The TSS score and chromatin fragment complexity were virtually identical between the two runs, confirming that the staining and barcoding of protein tags did not impact scATAC quality (**Fig. 1d** and **Extended Data Fig. 1f**). Reassuringly, projection of antibody tag counts on cell types resolved and annotated by their chromatin accessibility profiles shows expected patterns of expression of canonical cell type markers, including mutual exclusivity of CD4 and CD8 expression in T cells, CD16 in NK cells and a subset of monocytes, and CD14 in a non-overlapping set of monocytes (**Fig. 1e** and **Extended Data Fig. 1h**). Notably, in most cases the cluster specificity of antibody tag counts aligns with the chromatin activity score of the corresponding gene locus, with markedly increased sensitivity (**Fig. 1e** and **Extended Data Fig. 1h,i,j**). As fixation and mild permeabilization prior to droplet-based scATAC-seq retain mitochondria^14^, we further recover 31% mitochondrial reads in this experiment, allowing us to profile mtDNA mutations jointly with protein levels and chromatin accessibility in single cells (**Fig. 1f**).

Finally, we further expand the utility of ASAP-seq by incorporating Cell Hashing^17, 18^. In Cell Hashing^17, 18^ and related methods^19–22^, sample multiplexing is enabled by barcoded oligo tags (hashtags) that are attached by a variety of means to all cells of a specific sample. Joint barcoding of hashtags with the cell’s transcriptome, reveals both the sample of origin for individual cells and the presence of cross-sample multiplets (with >1 hashtag above threshold).

TSA hashing reagents are compatible with the ASAP-seq bridging strategy, and are barcoded in droplets together with protein tags and accessible chromatin fragments, before recovery using a separate PCR indexing strategy (**Methods**). To demonstrate sample multiplexing and doublet detection (enabling overloading and thus more efficient experiments), we stained PBMCs with 4 TSA hashing antibodies (**Supplementary Table 1, tab ’Hashing’**) and recovered 13,772 cells that were successfully demultiplexed in 4 distinct populations, with 1,396 detected doublets, consistent with the expected number of doublets of 1,138 derived from a Poisson-based model (**Extended Data Fig. 1g**).

### ASAP-seq is a modular toolkit that enables sensitive protein capture irrespective of antibody conjugate or lysis conditions used

We next determined if UBIs, used in TSA family antibody:oligo conjugates, perform comparably to UMIs (in TSB products). UBIs are copied off of the bridge oligos to acquire a near-unique sequence string for counting purposes. We designed 10 nt UBIs with complexity approaching or exceeding the UMI complexity commonly used in scRNA-seq^23, 24^, but note that the length and complexity of UBIs can be altered for different applications. To formally compare UMI vs. UBI quantification of protein tags, we simultaneously co-stained PBMCs with a 1:1 ratio of TSA (UBI-based) and TSB (UMI-based) TBNK panel (**Fig. 2a**, **Supplementary Table 1, tab ’TBNK’**). During the barcoding step, both bridge oligos (BOA and BOB) were added to the reaction mix in equal concentrations to bridge their corresponding tags (bridging schemes shown in **Extended Data Fig. 1b** and **Extended Data Fig. 2a**). UBI-collapsed TSA counts show good correlation with UMI collapsed TSB counts across all 9 antibodies (Pearson’s *r*=0.44-0.93, depending on antibody), suggesting that the UBI can provide a reliable proxy for a UMI (**Fig. 2b**).

While ASAP-seq directly extends mtscATAC-seq, yielding comparable retention of mtDNA reads, we asked whether protein abundance and accessible chromatin could be robustly measured without concomitant mtDNA enrichment, which may be preferred in specific tissue types or experimental settings. To this end, we compared the original OMNI-ATAC-seq lysis protocol previously shown to deplete mtDNA^25^, and currently recommended for the 10x Genomics scATAC-seq assay (**Methods**), to the effects of lysis conditions for mtscATAC-seq^14^.

We fixed PBMCs stained with the TBNK panel (**Fig. 2a**), split them in two aliquots, and lysed with the mild mtscATAC-seq conditions described thus far (referred to as low loss lysis, LLL) or with the stronger OMNI conditions including digitonin and Tween-20 in the lysis buffer. While the lysis detergent composition had a dramatic effect on mtDNA retention (-18x drop in median mtDNA fragments per cell; **Fig. 2c**), it had little to no effect on the distribution of UBI or UMI tag counts (**Fig. 2d** and **Extended Data Fig. 2c**). Moreover, The correlations between UBI- and UMI- collapsed tag counts under stronger permeabilization are comparable to those in milder lysis conditions (**Extended Data Fig. 2b**, Pearson’s *r*=0.38-0.96), albeit with slight improvement for most antibodies. Overall, we conclude that the cell surface marker retention in the ASAP-seq workflow allows reliable measurement, irrespective of antibody:oligo reagent type or lysis conditions used, and without compromising ATAC-seq data (**Extended Data Fig. 2d,e,f**).

### ASAP-seq is compatible with detection of intracellular proteins

We hypothesized that the fixation and permeabilization steps inherent in the ASAP-seq workflow would provide an opportunity to also detect intracellular epitopes, which have been previously inaccessible in high-throughput methods combining protein detection with scRNA-seq^5–7^. To examine this, we stained PBMCs in two steps, for which we used different conjugate families for extracellular and intracellular markers to allow independent amplification of the two tag libraries and tuning of sequencing depth for the two classes of proteins in the event of differences in tag recovery. We labeled cells with the TSA TBNK panel comprising extracellular surface markers, followed by fixation, permeabilization and staining with three TSB antibodies directed against intracellular epitopes, ZAP70, Perforin (*PRF1*), and Granzyme B (*GZMB*), before transposition and barcoding (**Fig. 3a**, **Supplementary Table 1, tab ’intracellular’**). Accessible chromatin profile-based clustering (**Fig. 3b**) and distribution of protein tags for extracellular markers within these clusters (**Fig. 3c**) was consistent with previous experiments and corresponding gene activity scores (**Extended Data Fig. 1i,j** and **Extended Data Fig. 3a,b**), verifying that the detection of these modalities remains robust to the additional intracellular staining step.

**Figure 3.**
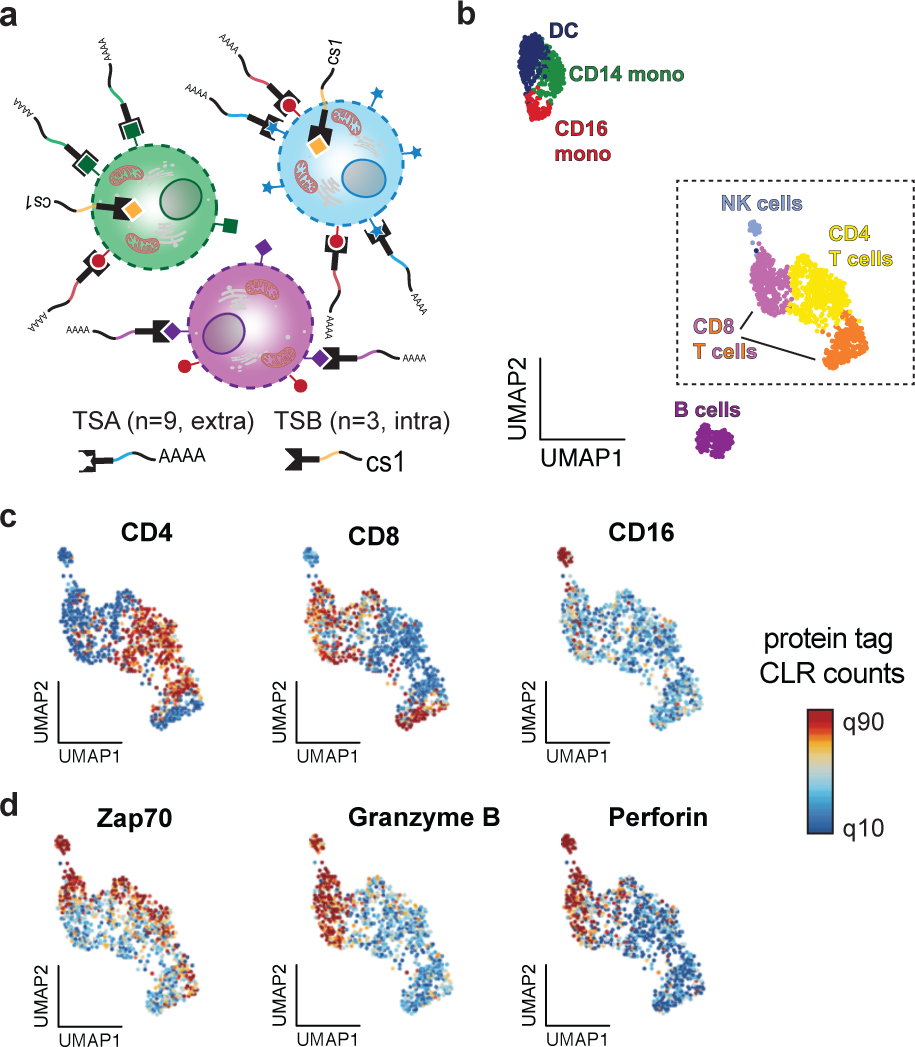
ASAP-seq enables detection of intracellular proteins with barcoded antibodies. a. Schematic of the intracellular staining experimental design. PBMCs stained with the TSA TBNK panel directed against cell surface markers were fixed, lysed, and stained with TSB antibodies directed against intracellular markers, followed by transposition. **b.** Two-dimensional embedding of the PBMC chromatin accessibility data using UMAP, with major peripheral blood cell types highlighted. **c,d.** T cells and NK cells as highlighted in the dashed-line box from panel (**b**) with superimposed tag intensities for indicated (**c**) cell surface and (**d**) intracellular markers. Color bar: protein tag CLR values.

Examining the distribution of protein tags for the intracellular proteins, we observed consistent expression as expected in the corresponding cell populations, with ZAP70 present in activated NK and T cells (CD4 and CD8), and Perforin and Granzyme B most prominent in natural killer (NK) cells and a subset of cytotoxic CD8^+^ T cells (**Fig. 3c,d**). Our observed intracellular abundances were indeed cell-type specific and further correlated with gene activity scores, ultimately validating the on-target activity for all three tested intracellular markers (**Fig. 3c,d** and **Extended Data Fig. 3a,b**). Thus, ASAP-seq enables the precise quantification of both cell surface and intracellular proteins, enabling new possibilities of defining cell states in single-cell genomics assays.

### ASAP-seq reveals cell state and cell lineage in human bone marrow

The multimodal readout of ASAP-seq uniquely enables profiling of epigenomic, proteomic, and clonal features (through mtDNA) of cells from native human tissue in a high-throughput manner. We applied ASAP-seq to profile bone marrow mononuclear cells from a healthy 24-year old donor, using a TSA antibody panel (n=242 markers, **Supplementary Table 1, tab ’BM’**) and 6 hashing antibodies to increase cell throughput. We permeabilized cells under LLL conditions to retain mtDNA fragments and barcoded in parallel with accessible chromatin fragments and protein tags (**Fig. 4a**). We retained 10,928 high-quality cells based on a combination of protein and chromatin-based quality control metrics (**Fig. 4a; Methods**). Dimensionality reduction and clustering based on single-cell chromatin accessibility profiles partitioned the cells into 21 distinct clusters spanning the major hematopoietic lineages, including progenitor and more differentiated lymphoid and myeloid cells (**Fig. 4b**; **Supplementary Table 2**). Notably, we did not remove predicted cell doublets as defined based on cellular markers, because these were enriched for monocytic progenitors, a cell state/type which was present in the donor’s bone marrow at the expected frequencies (**Extended Data Fig. 4a,b**; **Methods**) and because there was no support to remove them from hashtag collisions. This result reinforces the utilization of orthogonal technologies to detect cell doublets, such as hashtag antibodies shown here, to limit erroneous inferences about real or synthetic cell populations in complex tissue types, while simultaneously increasing throughput.

**Figure 4.**
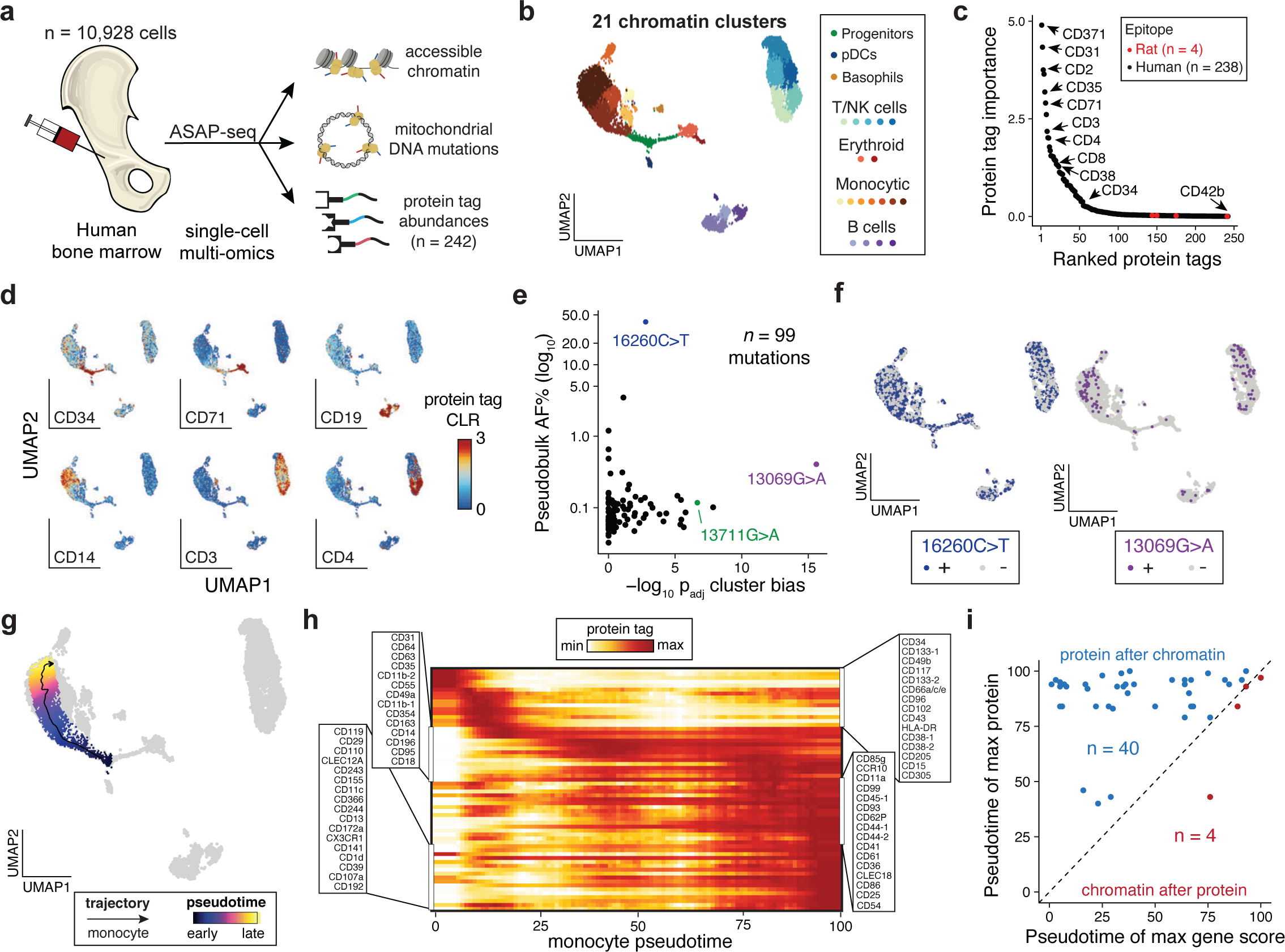
Dissection of native human hematopoiesis with multi-modal cell state inference and mtDNA-based lineage tracing. a. Schematic of experimental design. Whole human bone marrow mononuclear cells (BMMCs) were stained with hashtag antibodies and a 242 antibody panel for ASAP-seq processing. **b.** Reduced dimension representation and cell clustering of high-quality cells (n=10,928) inferred using chromatin accessibility. **c.** Rank sorting of informative protein tags in distinguishing cell cluster identification. Negative controls (rat epitopes) are shown in red. **d.** Characterization of cell populations for 6 selected markers. **e.** Characterization of 99 somatic mtDNA mutations identified in the BMMCs. Selected mutations enriched for lineage bias (13069G>A and 13711G>A; x-axis; see **Extended Data** Figure 4e) and highest for allele frequency (16260C>T; y-axis) are highlighted. **f.** Projection of highlighted mutations from (**e**) on the reduced dimension space. Thresholds for + were 50% for 16260C>T and 5% for 13069G>A based on empirical density. **g.** Developmental trajectory of monocyte differentiation using semi-supervised pseudotime analysis. **h.** Expression of cell surface markers along the developmental trajectory highlighted in (**g**). Rows are min-max normalized. **i.** Comparison of maximum gene activity scores (x-axis) and protein (y-axis) during pseudotime. Each dot is a gene/surface protein pair.

To identify protein markers associated with chromatin accessibility-derived cell subsets, we utilized and interpreted a Random Forest model trained on cell cluster labels using the scaled antibody tag abundances as previously implemented in CiteFuse^26^ (**Methods**). The model automatically rediscovered many widely-used surface markers for discriminating cell types in hematopoietic lineages, confirming its validity, including CD3, CD4, and CD8 in lymphoid cells, CD371 (*CLEC12A*) and CD2 in myeloid cells, CD71 (*TFRC*) in erythroid cells, and CD38 in more mature progenitor cells (**Fig. 4c,d**; **Extended Data** **Fig. 4c**). These analyses confirm that ASAP-seq can correctly uncover key surface proteins delineating cell types in complex tissue.

Next, we used the concomitant measurement of mtDNA genotypes for the clonal tracing of native hematopoietic cells^14, 27, 28^ within the human bone marrow compartment. Using mgatk^14^ we detected 99 heteroplasmic mtDNA mutations, which were enriched for classes of nucleotide substitutions consistent with previous reports^14^ (**Extended Data Fig. 4d**). Hypothesizing that some somatic mutations may be clonally associated with a specific lineage, we utilized cell subset annotations to examine for such putative (lineage) bias, thereby revealing insights into the functional heterogeneity of hematopoietic clones during blood production (**Fig. 4e; Methods**). Interestingly, somatic mutations such as 13069G>A and 13711G>A were relatively depleted in cells from the erythroid lineage (**Fig. 4e,f** and **Extended Data Fig. 4e,f**). Functional annotation of these mutations showed no predicted loss or gain of function, suggesting these somatic mtDNA mutations may mark lineage-restricted clones. Furthermore, one highly heteroplasmic variant, 16260C>T, was present at -40% heteroplasmy in the population, more than tenfold greater than any other detected somatic mutation, and yet was evenly distributed across the different hematopoietic lineages (**Fig. 4e,f**). Analysis of the donor mitochondrial haplotype suggested that this mutation indeed arose somatically, potentially presenting an early event during developmental hematopoiesis^29^ or alternatively, representing a population that clonally expanded during adulthood. While further studies will be needed to identify the molecular drivers underlying these dynamics, our observations support the utility of ASAP-seq to uncover somatic mtDNA variants and putative functional features of human hematopoiesis at single cell resolution.

### Dynamics of surface proteins during differentiation

While distinct single cell genomic measurements have revealed the continuous nature of hematopoietic differentiation^30, 31^, we hypothesized that the integration of accessible chromatin and protein tags via ASAP-seq could highlight an additional and distinct layer of surface protein marker dynamics during lineage commitment and differentiation, a process that has been traditionally characterized using a more limited set of markers. Utilizing a semi-supervised pseudotime approach, we charted trajectories from CD34+CD38- multipotential hematopoietic stem and progenitor cells (HSPCs) to differentiated monocytes (**Fig. 4g**) and erythroblasts (**Extended Data Fig. 4g**). While the protein expression of markers associated with multi-potent and other lineage progenitor cells, such as CD34 and CD49d (*ITGA4*), was down-regulated early in the trajectory, monocyte markers such as CD64 (*FCGR1A*) and CD31 (*PECAM1*) were quickly upregulated and persisted throughout differentiation (**Fig. 4h**; **Supplementary Table 3**). Conversely, markers such as CD11c (*ITGAX*) and CD371 (*CLEC12A*) were only upregulated toward the end of the trajectory. Though limited by cell number, we observed similar patterns of dynamic surface marker expression throughout erythroid differentiation (**Extended Data Fig. 4h**). These results demonstrate that ASAP-seq, as a multimodal assay with a large number of measured markers, can provide a substantially deeper profile of cell marker diversity in complex tissues than conventional flow and mass cytometry approaches^32^.

As ASAP-seq concomitantly measured accessible chromatin for these cells, including at the promoters of genes encoding these surface markers, we sought to examine how these modalities may be intertwined during differentiation. Among the proteins that were gained after commitment from the progenitor cluster, in the vast majority of cases, the increase in expression during monocyte differentiation was preceded by a gain of accessible chromatin at associated loci (**Fig. 4i** and **Extended Data Fig. 4i; Methods**). A similar pattern was observed, albeit with fewer markers, in erythroid differentiation (**Extended Data Fig. 4h,j**). This result is consistent with a model where chromatin accessibility is the ’first mover’ during differentiation and the resultant changes in transcription prime cells for differentiation^8^. Ultimately, we note that the disparity between binary chromatin accessibility versus accumulation of protein for single cells requires careful consideration of these modalities for understanding regulatory models. Taken together, our analyses showcase the versatility of ASAP-seq to measure multiple modalities of cell state alongside cell lineage, enabling an additional and distinct tool in the study of complex human tissues.

### ASAP-seq and CITE-seq reveal three levels of genetic regulation following stimulation

ASAP-seq and CITE-seq are companion assays that profile the epigenomic or transcriptional landscapes of single cells, respectively, together with the same highly multiplexed protein measurements. We reasoned that the shared protein features can help connect scRNA-seq and scATAC-seq datasets.

We thus applied both ASAP-seq and CITE-seq to profile epigenomic, transcriptomic, and proteomic changes following T cell stimulation. We split a single PBMC sample into two aliquots, one stimulated with tetrameric anti-CD3/CD28 in the presence of IL-2 for 16 hrs, while the other was cultured in the absence of stimulation (control) (**Methods**), followed by staining with a TSA antibody panel of n=227 antibodies (**Supplementary Table 1, tab ’PBMC’)**. Each of the samples was then split to run ASAP-seq and CITE-seq in parallel (**Fig. 5a**). We combined both the RNA and ATAC profiles from the control and stimulated cells, primarily revealing stimulation-dependent changes within the T cell population (**Fig. 5b,c**; **Methods**).

**Figure 5.**
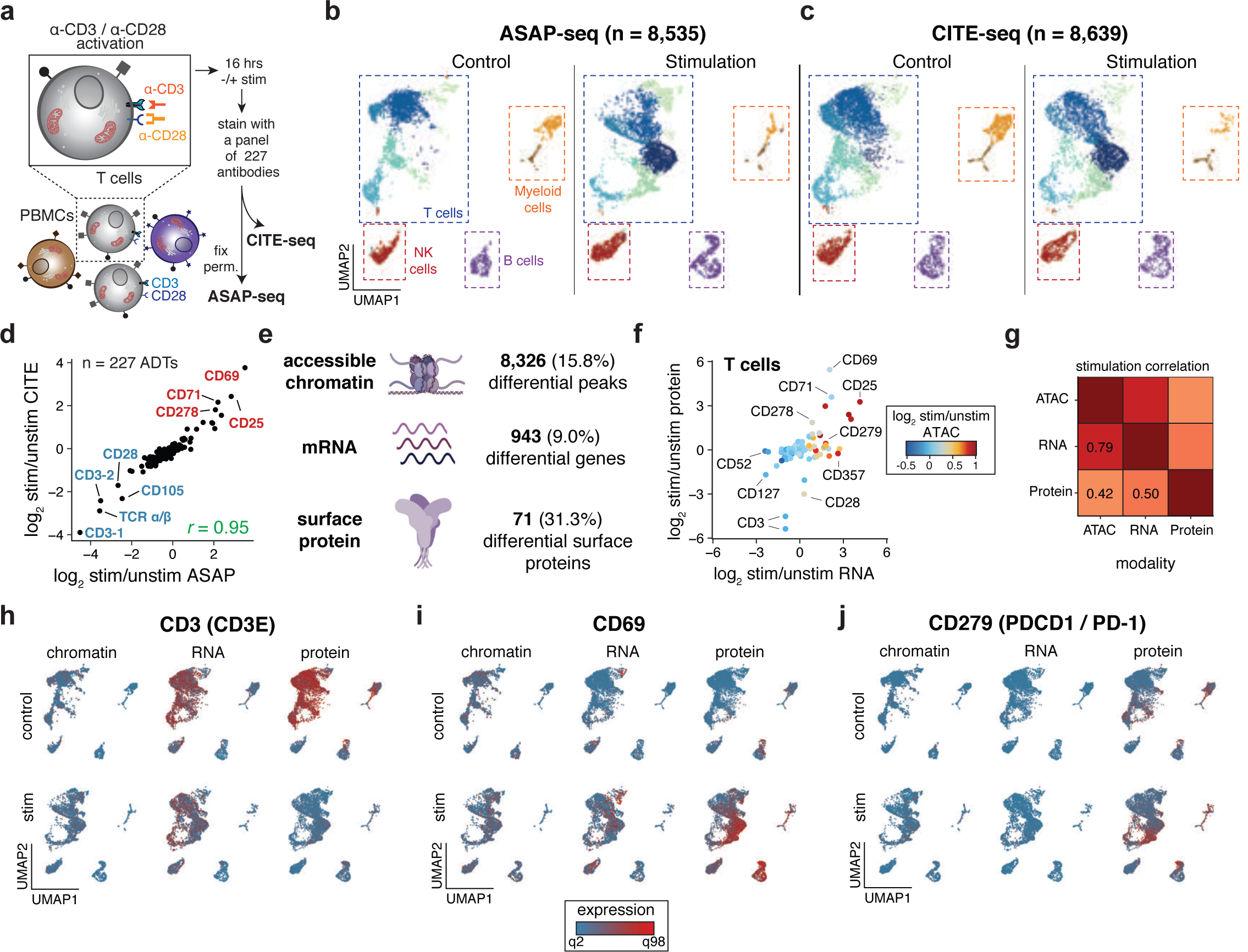
ASAP-seq and CITE-seq reveal coordinated and distinct changes in chromatin, RNA, and protein levels. a. Schematic of the experimental design. PBMCs were incubated with (stimulation) or without (control) multimeric a-CD3/CD28 for 16 hrs, followed by staining with the 227 antibody panel. An aliquot of the cells was withdrawn and subjected to CITE-seq, while the remaining cells were fixed and subjected to ASAP-seq. **b,c.** Reduced dimension representations using data integration methods and UMAP for (**b**) ASAP-seq and (**c**) CITE-seq for both control (left) and stimulated conditions (right). **d.** Correlation of surface marker fold changes (log_2_) upon stimulation as detected by CITE-seq and ASAP-seq. Top upregulated markers are highlighted in red, and downregulated in blue. **e.** Schematic and summary of number and proportion of differential features (chromatin accessibility peaks, genes, and surface proteins) detected for T cells between the stimulation and control. **f.** Summary of changes in chromatin accessibility, gene expression and surface protein abundance for 84 expressed genes during T cell stimulation. **g.** Pearson correlation between the log_2_ fold changes for each modality as shown in (**f**). **h-j.** UMAPs of single-cell chromatin accessibility, mRNA expression, and surface protein levels for both the control (top) and stimulation condition (bottom) shown on the reduced dimension space for (**h**) CD3, (**i**) CD69 and (**j**) PD-1.

As protein abundances were determined in the same population of cells, we directly compared surface protein measurements inferred by CITE-seq and ASAP-seq. As expected, we observed a decrease (-1.7-2x) in the tag molecule complexity in ASAP-seq compared to CITE-seq, consistent with the drop in fluorescence intensity observed by flow cytometry due to the additional processing necessary in ASAP-seq (**Extended Data Fig. 5a, 1a; Methods**). However, the two methods were highly concordant in the change in antibody signal stimulation across the panel (Pearson’s *r*=0.95, **Fig. 5d**). Importantly, we did not observe specific loss of any markers in ASAP-seq relative to CITE-seq, indicating that the cell processing-induced loss of sensitivity is a general phenomenon that does not specifically affect a subset of markers. Notably, both assays detected substantial upregulation of canonical T cell activation marks, such as CD69, CD25, CD71 (*TFRC*), and CD278 (*ICOS*)^33–35^, at both the pseudobulk (**Fig. 5d**) and single-cell (**Extended Data Fig. 5b,c**) level. Conversely, CD3 (*CD3E*; protein log_2_FC=-3.5 and -4.5; p<2.2×10^-16^, Wilcoxon rank sum test for ASAP-seq protein abundance), CD28 (log_2_FC=-2.5 p<2.2×10 ^-16^), and TCR α/β (log FC=-2.9; p<2.2×10 ^-16^) antibody counts were noticeably reduced upon stimulation (**Fig. 5d**), likely due to internalization of the engaged and non-engaged receptors upon triggering of the TCR complex^36^. An antibody prioritization approach utilizing the Random Forest model^26^ for ASAP-seq data (**Methods**) verified that these markers were most associated with the stimulation at single cell resolution (**Extended Data Fig. 5d**). Notably, other canonical lineage markers, such as CD4 and CD56 (*NCAM1*), were prioritized in distinguishing cell states inferred by chromatin accessibility clustering irrespective of stimulation. Finally, embedding cells by protein abundance profiles intermixed cells profiled by the two assays, albeit with reduced separation of the activated T cell state, likely due to the relatively modest number of dynamically responding proteins compared to accessibility peaks or transcripts (**Fig. 5e** and **Extended Data Fig. 5e-g**). Taken together, these analyses and results indicate that despite a lower tag complexity, ASAP-seq is similarly capable of capturing protein abundance associated with cell state and dynamic changes as measured with CITE-seq.

To characterize the overall cellular response to stimulation, we examined the dynamic changes in accessible chromatin, gene expression, and protein abundance in stimulated vs. control T cells. At consistent magnitude and statistical significance thresholds, we detected 8,326 differential peaks, 943 differentially expressed genes, and 71 differentially expressed surface proteins, consistent with previous unimodal analyses largely from bulk experiments^37, 38^ (**Fig. 5e; Methods**). Of the 84 cases where all three modalities were detected in T cells, we observed heterogeneous responses in gene expression, chromatin accessibility, and surface protein abundance (**Fig. 5f**; **Supplementary Table 4**), with chromatin and protein changes being the least concordant (**Fig. 5g**). Specifically, CD3 (*CD3E*) and CD28 downregulation along with CD69 upregulation are striking on the protein level, evident transcriptomically only for *CD3E* and *CD69*, but barely detectable at the chromatin accessibility level (**Fig. 5h,i** and **Extended Data Fig. 5h**). This can be due to true invariance in chromatin accessibility, such that gene expression is temporarily repressed without loss of accessibility, or to technical challenges, for example given the higher sensitivity in capturing a modality with higher copy number (protein), as exemplified by CD4 and CD279 (*PDCD1* or *PD-1*) (**Fig. 5j** and **Extended Data Fig. 5i**). On the other hand, we observed RNA-specific changes in CD52 where chromatin accessibility and protein abundance were relatively constant pre/post stimulation (**Extended Data Fig. 5j**). Together, these analyses and anecdotes highlight the utility of combining ASAP-seq and CITE-seq to distinguish changes at three levels of gene regulation.

Because we activated T cells in a multicellular PBMC culture, we next leveraged the single-cell nature of our data to identify secondary effects in other cell subsets. In particular, we examined corresponding changes across the three modalities in B cells, focusing on a set of 103 well-expressed proteins (**Extended Data Fig. 5k**). While many changes mirrored those of T cells (likely due to low-frequency doublets in our annotated B cell clusters; **Methods**), we highlight two markers, CD25 (*IL2RA*) and CD184 (*CXCR4*), where RNA and protein, but less notably chromatin accessibility, were affected, either in direct response to IL2 or as a secondary response to T cell stimulation (**Extended Data Fig. 5l,m**). Notably, CD25^+^ B cells have been reported to possess enhanced antigen presentation capabilities, which may in turn facilitate enhanced T cell responses^39^. Furthermore, prior work has indicated that IL-21 (produced by activated T cells) accelerates CXCR4 internalization in B cells, which may be important for the regulation of B cell homeostasis^40, 41^. As the loss of CXCR4 expression has traditionally been observed in germinal centers^40, 41^, our inclusive antibody panels enable the observation of such changes across a multitude of cell states.

Taken together, our CD3/CD28 stimulation system directly highlights how the single-cell, three-tier characterization by ASAP-seq and CITE-seq can dissect correlated and discordant changes in complex tissue settings.

### Multiplexed CRISPR perturbations in primary T cells

As ASAP-seq revealed distinct chromatin and protein changes underlying normal T cell activation, we sought to refine some of the potential underlying mechanisms by targeted perturbations via an arrayed CRISPR/Cas9 screening strategy in primary human cells. To this end, we purified naive human CD4^+^ T cells from peripheral blood of three healthy donors, pooled and stimulated them with anti-CD3/CD28 beads for 72 hours. After four days of resting, we transfected cells by electroporation with Cas9 protein individually complexed with gRNAs targeting *CD4, ZAP70, NFKB2* (2 gRNAs per gene), *CD3E*, *CD3E+CD4* (dKO), or one of two non-targeting controls (NTCs; **Fig. 6a**)^42, 43^. Following electroporation, cells were rested for an additional 7 days in the presence of IL-2 before re-stimulation with anti-CD3/CD28 beads for another 72 hours (**Extended Data Fig. 6a**). We multiplexed each of the ten perturbation conditions post-stimulation using a unique combination of two TSA hashing antibodies as a surrogate for gRNA identities and then stained the cells with an antibody panel (n = 37) before downstream processing by the ASAP-seq workflow (**Supplementary Table 1, tabs ’hashing’ and ’perturbation’)** (**Supplementary Table 5**). Demultiplexing by hashtag reads from perturbed cells enabled high-confidence identification of 5,825 single perturbed cells approximately evenly distributed across all perturbation conditions with a median yield of 1.47 × 10^4^ fragments mapping to the nuclear genome (**Extended Data Fig. 6b**).

**Figure 6.**
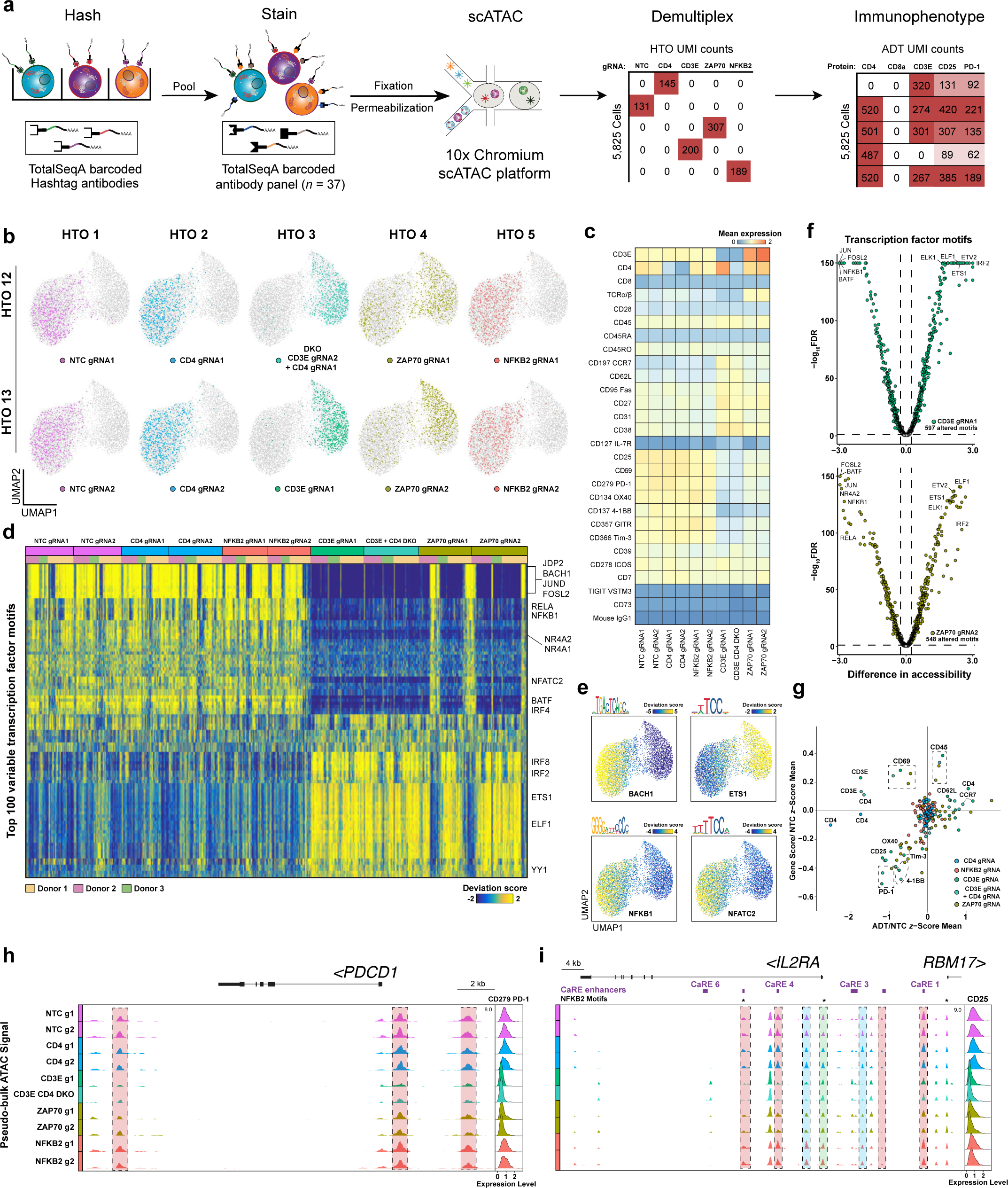
Multiplexed CRISPR perturbations with ASAP-seq in primary human T cells. a. Schematic workflow for combinatorial multiplexing with ASAP-seq. CRISPR-edited cells are first stained with oligo-conjugated hashtag antibodies and then pooled for downstream processing by ASAP-seq. gRNA identities are demultiplexed using hashing antibody counts. **b.** UMAP embedding of *n* = 5,825 single cells and their associated gRNAs. **c.** Heatmap showing mean expression for 27 selected surface protein markers across gRNA perturbations in stimulated cells. **d.** Heatmap representation of chromVAR bias-corrected transcription factor motif deviation scores for the top 100 most variable transcription factors across perturbation conditions. Associated gRNA and donor information are color-coded and indicated at the top of the plot. **e.** Overlay on ASAP-Seq UMAP of chromVAR transcription factor motif deviations. The motif for the given transcription factor is indicated at the top of the plot. **f.** Volcano plots showing transcription factor motifs with significantly changed chromatin accessibility profiles between NTC cells and guides targeting *CD3E* and *ZAP70* (FDR <= 0.05, chromVAR accessibility change >= 0.25). **g.** Scatterplot of mean gene activity scores for 22 individual gene loci plotted against CLR-normalized mean protein tag counts associated with each gRNA. Values are normalized against NTC cells. **h,i.** Genomic tracks of (**h**) *PDCD1* (gene encoding PD-1) and (**i**) *IL2RA* (gene encoding CD25), indicating pseudo-bulk ATAC signal tracks across gRNAs with corresponding CLR-normalized protein abundance ridge plots. Differentially accessible regions are highlighted in red. Differentially accessible regions not overlapping CARE enhancers are highlighted in blue (**i**) and the TSS is highlighted in green (**i**). NFKB2 sequence motif matches are indicated by *.

Cells perturbed by gRNAs targeting critical regulators of TCR signal transduction (*CD3E* and *ZAP70*) had similar chromatin accessibility profiles and clustered together (**Fig. 6b**). Moreover, these cells largely expressed protein markers characteristic of a resting state, such as CD197 (*CCR7*) and CD62L (*SELL*), indicating a profound defect in TCR stimulation response (**Fig. 6c** and **Extended Data Fig. 6d**). In contrast, cells with non-targeting (NTC) gRNAs or with gRNAs targeting *CD4* and *NFKB2* clustered together and displayed high levels of surface protein expression for classical T cell activation markers such as CD25, CD69, CD137 (*TNFRSF9*) and CD279 (*PDCD1* or *PD-1*), indicating active TCR signaling. Importantly, only cells with gRNAs targeting *CD4* exhibited substantial reduction in CD4 surface protein expression, further validating the robustness of the workflow and reliability of the assay (**Fig. 6c** and **Extended Data Fig. 6c**).

Next, we refined our findings from our PBMC stimulation experiment by assessing protein expression impacted by each gRNA perturbation upon re-stimulation. In line with our expectations, (**Fig. 5d,h**), ZAP70-deficient cells exhibited increased expression of TCRa/1 and CD3E (**Fig. 6c**), demonstrating that downstream TCR signaling is necessary to mediate downregulation of these molecules. TCRa/1 expression was similarly absent in *CD3E*-perturbed cells, in agreement with its reliance on the CD3 complex for proper surface localization^44^. By contrast, CD28 expression was only marginally detected across all perturbations, suggesting that CD28 signaling alone is sufficient for its internalization from the surface. Consistent with its known role as an inducible co-stimulatory molecule^45, 46^, we also found that CD278 (*ICOS*) expression could be promoted and sustained with intact CD28 signaling, independent of effective CD3E stimulation (**Fig. 6c**). Together, these analyses verify the utility of ASAP-seq in revealing how targeted gene manipulation affects protein expression changes in activated cell states.

We next inferred changes in gRNA-dependent transcription factor activities by quantifying accessible transcription factor motif deviations using chromVAR^47^. We found that gRNAs targeting the same gene had similar predicted effects, despite varying targeting efficiencies (revealed by discrepancies between hashtag-defined perturbation identities and cellular phenotypes, consistent with variable guide editing efficiencies as quantified by next-generation sequencing) and donor origin (**Fig. 6d-f** and **Extended Data Fig. 6e-g**). As expected, in comparison to NTC cells, depletion of *CD3E* resulted in a defective response to TCR re-stimulation and significantly decreased accessibility in regions containing motifs of Activator Protein-1 (AP-1) transcription factor family proteins such as c-JUN and BATF (median chromVAR accessibility loss, 10.24; FDR < 0.0001; chromVAR accessibility loss 6.96; FDR < 0.0001, respectively). Similarly, gRNAs targeting *ZAP70*, an immediate kinase effector of TCR signaling, displayed a modest decrease in AP-1 family motif accessibility, but the effect was more bimodal possibly due to a delayed effect or less efficient editing in some cells. Additional altered transcription factor motifs in *CD3E-* and *ZAP70*-targeted cells included NFAT family transcription factors, consistent with their crucial roles in chromatin remodeling and transcriptional regulation following TCR activation^48, 49^. Interestingly, disruption of *NFKB2* led to an increase of accessibility for NFKB family motifs, which could be reflective of competitive dimerization of p50 and p52 for common binding partners RELA and RELB (**Fig. 6d** and **Extended Data Fig. 6f**)^50^. These results demonstrate that the multi-modal ASAP-seq readouts can successfully resolve stimulation-responsive transcription factor-chromatin interactions in the context of genetic perturbations.

While an existing method that also pairs measurements of perturbations and ATAC-seq profiles in single cells has enabled the dissection of molecular machinery governing cell state^51^, this existing approach yields low cell numbers that limits observations. Conversely, our approach uniquely allows queries of how specific chromatin changes may relate to changes in protein expression. To examine this, we first compared accessible chromatin scores with concomitant surface protein profiling across each perturbation condition for 22 individual gene loci (**Supplementary Table 5**). Overall, perturbation-induced changes in surface protein expression were correlated with changes in chromatin status (*r* = 0.57 in surface proteins not targeted by a perturbation and *r* = 0.70 when further excluding CD69; **Fig. 6g,h**). For example, many stimulation-responsive genes such as CD25, CD134 (*TNFRSF4*), and CD279 (*PDCD1* or *PD-1*) were downregulated in both protein expression and chromatin accessibility in CD3E- and ZAP70-targeted cells (when compared to NTC cells). In contrast, CD69 gene accessibility was slightly increased with a significant decrease in surface protein levels. Overall, these dynamic changes in our perturbation system mirrored our results in our previous stimulation system (**Fig. 5**). Interestingly, we observed a more pronounced coordination between changes in protein expression and gene activity for CD357 (*TNFRSF18*) and CD366 (*HAVCR2*; CLR-normalized mean protein tag difference of 0.84 and 0.66, respectively, between CD3E-targeted cells and NTC; **Extended Data Fig. 6h**). This was not evident in our PBMC stimulation experiment, where changes in CD357 and CD366 protein levels were only modest, despite increased accessibility at associated stimulation-responsive enhancers (CLR-normalized mean protein tag difference of 0.18 and 0.17, respectively, between CD4 T cell stimulation and control), likely due to the shorter, 16-hour stimulation period.

Finally, as recent efforts in fine-mapping *cis*-regulatory elements by CRISPR screening have enhanced the capacity to uncover functional regulatory elements in different contexts^52–55^, we reasoned that our perturbation screening approach coupled with ASAP-seq could offer similar biological insights in identifying stimulation-responsive accessible chromatin regions. Examining pseudo-bulk gRNA-associated ATAC signal tracks at the *IL2RA* locus, we found strong depletion of chromatin accessibility in a number of regions with a concomitant decrease in the expression of CD25 (IL2RA) protein for cells targeted by gRNAs against *CD3E* and *ZAP70*, suggesting a prerequisite of TCR stimulation in the activation of these putative enhancers (**Fig. 6i**). These impacted enhancers largely overlapped the *IL2RA* CRISPRa-responsive elements (CaREs) that have been previously characterized by functional screening^54^. In particular, we observed marked accessibility changes overlapping CaRE4, which has been validated as a TCR stimulation-responsive enhancer for *IL2RA*. Conversely, CaRE3, which has recently been labeled as a Treg-specific enhancer, was indeed relatively static in our system^56^. Moreover, we observed a decrease in CD25 expression in cells perturbed by gRNAs targeting *NFKB2*, despite relatively unchanged chromatin accessibility and the presence of compatible NFKB2 DNA binding motifs within regulatory regions. These results suggest that while NFKB2 does not actively regulate local chromatin accessibility at this locus, it may still play a role in coordinating and maintaining CD25 expression in activated T cells. Taken together, our integrated approach enabled by the multi-modal readouts of ASAP-seq allows for unbiased discovery of context-dependent coding and non-coding gene regulation modules.

## DISCUSSION

Here, we present ASAP-seq, a unique approach that enables the concomitant detection of protein abundance alongside transposase-accessible chromatin and mtDNA in thousands of single cells. Our method is enabled by recent modifications to droplet-based scATAC-seq, notably the retention of the cellular membrane following pooled permeabilization, which further enables the simultaneous, efficient sequencing of mtDNA^14, 15^. As the majority of cell atlases to date have characterized the distinct transcriptomes of single cells in complex tissue, ASAP-seq provides a complementary multi-omic approach to map regulatory elements, protein abundances, and clonal relationships.

The ASAP-seq workflow is directly compatible with related multimodal assays that simultaneously measure protein and RNA, namely CITE-seq^5, 6^, by utilizing oligo:antibody conjugates to infer protein abundances in complex cell mixtures (**Fig. 5**). Importantly, our approach introduces a bridge oligo (**Fig. 1**) that enables the utilization of existing antibody conjugates, yielding an accessible and user-friendly protocol. As the multimodal toolkit continues to evolve, our bridge oligo innovation will provide important flexibility to append protein quantification to other assays, most immediately, transposon-based methods such as CUT&Tag^57^.

Furthermore, we show that our protocol is compatible with the direct detection and quantification of intracellular markers (**Fig. 3**). While other protocols have achieved concomitant quantification of intracellular protein abundance and gene expression with plate-based methods that also require specialized fixation conditions^58^, or combining FACS-based enrichment of cells with scRNA-seq^59^, ASAP-seq provides a more parsimonious approach to concomitant estimation of different marker types on a widely-used, high-throughput commercial platform. Moreover, protein detection and quantification remain paired with the chromatin accessibility profile of each cell, thereby reflecting the full dynamic range of protein expression. Thus, the combination of detecting extracellular and intracellular protein abundances, which we show can be encoded by different capture sequences, enables a distinct mechanism to chart cell states and their underlying regulatory elements. We anticipate the demonstration of intracellular protein detection by ASAP-seq will spur the development of large panels of oligo labeled antibodies targeting intracellular epitopes ranging from signaling molecules, specific phospho-epitopes, or to transcription factors.

By examining our multimodal readouts in native and perturbed hematopoietic tissue, our analyses reveal distinct cellular programming occurring in chromatin, transcriptional, and post-translational regulation. In particular, we observe chromatin-based priming during a monocyte developmental trajectory in the native bone marrow (**Fig. 4**). Conversely, during T cell activation, we observe a more heterogeneous response where changes in chromatin, RNA, and protein abundances become more uncoupled (**Fig. 5**). By further utilizing CRISPR-based perturbations (**Fig. 6**), we disentangle downstream signalling contributions of the TCR and CD28, providing a blueprint for the scalable study of combined cellular chromatin and protein expression dynamics in human cells. Furthermore, our analyses in the *IL2RA* locus reveal how ASAP-seq can enable the fine-mapping of regulatory elements in various cell states that directly impact protein expression. Future extensions of ASAP-seq that incorporate direct detection of guide sequences^6, 60^, or encoded guide barcodes^61–63^ will further enable pooled screens at a substantially increased scale.

We anticipate that ASAP-seq, when coupled with large antibody panels which exceed the antigen diversity measurable by current mass cytometry approaches^32^ (as demonstrated in this study), may facilitate the discovery of deregulated surface markers on (pre-)malignant/leukemic (stem) cell populations that could be further exploited for diagnostics or antibody-mediated therapies^64, 65^. Further, as showcased by the sensitivity of our measurement of PD-1 in activated T cells (**Fig. 5,6**), ASAP-seq should provide a powerful molecular approach to identify and study the epigenetic dysfunction in distinct states of adaptive immune cells in tumors, infectious diseases, and other malignancies. In total, our methodological approach and analyses indicate that ASAP-seq provides a modular, powerful toolkit for understanding the behavior of single cells in complex settings.

## ONLINE METHODS

See Protocol Exchange (doi pending) or CITE-seq.com/protocols for a step-by-step ASAP-seq protocol.

### Cells

Cryopreserved healthy donor peripheral blood mononuclear cells (PBMCs) and bone marrow cells (BM) were obtained from AllCells (USA) or Cellular Technology Limited (CTL) and processed immediately after thawing. NIH-3T3 and HEK293FT cells were maintained according to standard procedures in Dulbecco’s Modified Eagle’s Medium (Thermo Fisher, USA) supplemented with 10% fetal bovine serum (Thermo Fisher, USA), at 37°C with 5% CO2.

### Cell staining with barcoded antibodies

TSA and TSB conjugated antibodies and panels were obtained from Biolegend, see **Supplementary Table 1** for a list of antibodies, clones and barcodes used for ASAP-seq. Cells were stained with barcoded antibodies as previously described for CITE-seq^5, 6^. Briefly, approximately 1.5-2 million cells per sample were resuspended in 1x CITE-seq staining buffer (2% BSA, 0.01% Tween in PBS) and incubated for 10 min with Fc receptor block (TruStain FcX, BioLegend, USA) to block FC receptor-mediated binding. Subsequently, cells were incubated with indicated antibodies or panels for 30 min at 4°C, as recommended by the manufacturer (BioLegend, USA). After staining, cells were washed 3x by resuspension in 1x CITE-seq staining buffer followed by centrifugation (300 g, 5 min at 4°C) and supernatant exchange. After the final wash, cells were resuspended in PBS and subjected to fixation and permeabilization as described in the section *Cell fixation and permeabilization*.

Intracellular staining was performed in fixed and permeabilized cells that were resuspended in Intracellular Staining Buffer (Biolegend, custom part number 900002577), with the addition of TruStain FcX and True Stain Monocyte blocker as recommended by the manufacturer (BioLegend).

### Cell fixation and permeabilization

Cells were fixed in 1% formaldehyde (FA; ThermoFisher, no.28906) in PBS for 10 min at room temperature, quenched with glycine solution to a final concentration of 0.125 M before washing cells twice in PBS via centrifugation at 400 g, 5 min, 4°C. Cells were subsequently treated with the appropriate lysis buffer depending on downstream application. If mtDNA retention was desired, permeabilization was performed as described in mtscATAC-seq^14^ with 10 mM Tris-HCl pH 7.4, 10 mM NaCl, 3 mM MgCl_2_, 0.1% NP40, 1% BSA (referred to as low loss lysis conditions or LLL). When mtDNA depletion was desired, cells were lysed in 10 mM Tris-HCl pH 7.4, 10 mM NaCl, 3 mM MgCl_2_, 0.1% NP40, 0.1% Tween20, 0.01% Digitonin, 1% BSA (referred to as OMNI conditions). permeabilization was performed on ice, 3 min for primary cells and 5 min for cell lines, followed by adding 1 ml of chilled wash buffer (10 mM Tris-HCl pH 7.4, 10 mM NaCl, 3 mM MgCl_2_, 1% BSA) and inversion before centrifugation at 500 g, 5 min, 4°C. The supernatant was discarded and cells were diluted in 1x Diluted Nuclei buffer (10x Genomics) and filtered through a 40 µm Flowmi cell strainer before counting using Trypan Blue and a Countess II FL Automated Cell Counter.

### Transposition and barcoding

Cell were subsequently processed according to the Chromium Single Cell ATAC Solution user guide (Versions CG000168 Rev D for v1 and CG000209 Rev D for v1.1, 10x Genomics) with the following modifications:

1. During the barcoding reaction (step 2.1), 0.5 µl of 1 µM bridge oligo was added to the barcoding mix. The sequences of the bridge oligos are: BOA (bridge oligo for TotalSeq^TM^-A): TCGTCGGCAGCGTCAGATGTGTATAAGAGACAGNNNNNNNNNVTTTTTTTTTTTTTTTTTTTT TTTTTTTTTT/3InvdT/ and BOB (bridge oligo for TotalSeq^TM^-B): TCGTCGGCAGCGTCAGATGTGTATAAGAGACAGTTGCTAGGACCGGCCTTAAAGC/3InvdT/
2. To facilitate bridge oligo annealing during GEM incubation (step 2.5), a 5 min incubation at 40°C was added at the beginning of the amplification protocol (40°C 5 min, 72°C 5 min, 98°C 30 sec, 12 cycles of 98°C 10 sec, 59°C 30sec, 72°C 1 min, ending with hold at 15°C). This extra annealing step was not essential when using TSA products, but increased efficiency in TSB tag capture.
3. During silane bead elution (step 3.1o), beads were eluted in 43.5 µl of Elution Solution I and 3 µl were kept aside to use as input in the tag library PCR, while the remaining 40 µl were used to proceed with SPRI clean up as the protocol describes. We reasoned that some tag fragments could stay in the bound fraction during the 1.2x SPRI separation, so to maximize tag capture we recommend to include a small portion (up to 10%) of the silane bead elution as input in the tag indexing reaction.

During SPRI cleanup (step 3.2d), the supernatant was saved and an additional 0.8x reaction volume of SPRI beads (32 µl) was added to bring the ratio up to 2.0x. Beads were washed twice with 80% ethanol and eluted in EB. This fraction can be combined with the few µl left aside after the silane purification to be used as input in the protein tag indexing reaction, or either source can be used alone with minimal impact on tag complexity (see **Extended Data Fig. 1**). PCR reactions were set up to generate the protein-tag library (P5 and RPI-x primers for TSA conjugates, P5 and D7xx_s for TSB conjugates) and the hashtag library (P5 and D7xx_s) with the program: 95°C 3min, 14-16 cycles of 95°C 20sec, 60°C 30sec, 72°C 20sec, followed by 72°C for 5min and ending with hold at 4°C. Example of an RPI-x primer (TruSeq Small RNA handle, present in TSA tags. “x” nucleotides present a are user-defined sample index): CAAGCAGAAGACGGCATACGAGATxxxxxxxxGTGACTGGAGTTCCTTGGCACCCGAGAATT CCA. Example of an D7xx_s primer (TruSeq DNA handle, present in TSB tags or TSA hashing): CAAGCAGAAGACGGCATACGAGATxxxxxxxxGTGACTGGAGTTCAGACGTGTGC. The final libraries were quantified using a Qubit dsDNA HS Assay kit (Invitrogen) and a High Sensitivity DNA chip run on a Bioanalyzer 2100 system (Agilent).

Note: Both v1 and v1.1 versions of the scATAC kit were successfully used throughout this study, with no discernible differences with respect to protein tag detection.

### Flow cytometry

For flow cytometry analysis, PBMCs were washed in FACS buffer (1% FBS in PBS) before antibody staining using a FITC-conjugated CD19 antibody (HIB19, 302206, Biolegend) and a Pacific-Blue-conjugated CD3 antibody (HIT3a, 300330, Biolegend), each at a 1:25 dilution. After washing, fixation and permeabilization were conducted as described in the section Cell fixation and permeabilization above, before cells were resuspended in nuclei dilution and ATAC buffer and incubated at 1h, 37°C in a thermocycler to mimic the Tn5 transposition step during (mt)scATAC-seq. Aliquots for flow cytometry analysis were processed at indicated stages as schematically depicted in **Extended Data Fig. 1a**. Analysis was conducted on a BD Bioscience Fortessa flow cytometer at the Whitehead Institute Flow Cytometry core. Data was analyzed using FlowJo software v10.4.2.

### PBMC stimulation

Cryopreserved PBMCs were thawed and washed in complete medium (RPMI Glutamax, supplemented with 10% FCS and 50 IU/ml IL-2). Cells were allowed to rest in complete medium for 30 min at 37°C, before filtering through a 70 µm cell strainer to remove aggregates. PBMC aliquots were split in half and resuspended to a final density of 1×10^6^/ml in either complete medium (unstimulated control) or complete medium supplemented with ImmunoCult Human CD3/CD28 T cell Activator (stimulated sample) according to the manufacturer (StemCell Technologies). 200 µl cell suspension aliquots were deposited in a 96-well round bottom plate and placed in a humidified 5% CO2 incubator at 37°C, for 16 hrs. Cells from respective wells were pooled, harvested, washed 2x with media and resuspended in 1 ml media, before filtering through 70 µm to remove cell aggregates. About 1×10^6^ cells of each condition were then harvested and resuspended in 100 µl CITE-seq staining buffer in preparation for staining.

### Arrayed Cas9 Ribonucleotide Protein (RNP) preparation and electroporation

Lyophilized crRNAs and tracrRNAs (IDT) were reconstituted to a concentration of 400 µM and stored in -80°C until use. crRNAs and tracrRNAs were mixed at a 1:1 v/v ratio, transferred into a 96-well plate and heated at 95°C for 5 min, followed by incubation at room temperature for 15 minutes to complex the gRNAs. 30 µg Cas9 protein (TakaraBio, Cat# Z2640N) was added to each well and mixed by gentle pipetting, followed by incubation at room temperature for 15 minutes. Complexed RNPs were then dispensed in a 96-well V-bottom plate at 12.7 µL per well. Cells were resuspended in Lonza P2 primary nucleofection buffer at 1×10^6^ cells per 20 µL and added to the RNP-containing V-bottom plate. The mixture was gently mixed by pipetting and then transferred into a 16-well electroporation cuvette plate (Lonza, Cat# V4XP-2032) and pulsed with the EH100 program. Immediately following electroporation, 100 µL pre-warmed T cell culture medium was gently added to each well and cells were incubated at 37°C for 10 minutes. Cells were then transferred into 96-well U-bottom plates for culture at 1×10^6^ cells/ml, supplemented with 500 IU/ml IL-2. A list of all crRNAs used in this study can be found in **Supplementary Table 5**.

### Multiplexed perturbation workflow

Primary human CD4+ T cells were enriched by magnetic negative selection using the human CD4+ T cell Isolation Kit (Miltenyi, Cat# 130-096-533) as per manufacturer’s instructions. Cells were then stained and na“ve CD4 T cells were sorted on a BD FACSAria-SORP system (Becton Dickinson) on the basis of CD4 and CD45RA expression. After isolation, cells were cultured in T cell culture medium consisting of RPMI with 10% Fetal Bovine Serum, 10 mM HEPES, 2 mM GlutaMax (Gibco, Cat# 35050-061), 1x MEM Non-Essential Amino Acids (Gibco, Cat# 11140-050), 1 mM Sodium pyruvate, 55 µM 2-mercaptoethanol and 100 IU/ml IL-2 at a density of 1×10^6^ cells/ml, and stimulated with anti-human CD3/CD28 Dynabeads (ThermoFisher, Cat#11131D) at a 1:1 cells-to-beads ratio. 72 hours after stimulation (Day 3), beads were removed and cells were rested in media containing IL-2 for expansion, while maintaining at a density of 1×10^6^ cells/ml. On Day 7, cells were electroporated with Cas9 ribonucleoprotein (RNP) complexes. Following electroporation, cells were cultured in media with 500 IU/ml IL-2 and split regularly to maintain a density of 1×10^6^ cells/ml. On day 15, cells we re-stimulated with anti-human CD3/CD28 Dynabeads (ThermoFisher, Cat# 11132D), supplemented with 100 IU/ml IL-2. 72 hours later, beads were removed and cells for each condition were stained and washed as described above with a combination of two specific TotalSeqA hashtag antibodies (0.25 µg per antibody). Live cells were enriched and pooled by cell sorting on a BD FACSAria-SORP (Becton Dickinson) and then processed as per the ASAP-seq protocol described above using OMNI lysis conditions.

### Next-generation sequencing of DNA amplicons

Next-generation sequencing of gDNA was performed essentially as previously described^66^. Cells transfected with Cas9 were harvested eight days post-electroporation, enriched for live cells by cell sorting on a BD FACSAria-SORP (Becton Dickinson) and then processed for gDNA extraction using the DNeasy Blood & Tissue Kit (Qiagen, Cat# 69504) following the manufacturer’s instructions. Genomic sites of interest were first amplified by PCR with Phusion high-fidelity DNA polymerase (NEB) using gene-specific primers (primer sequences are listed in **Supplementary Table 5**). A second round of PCR was performed using 1 µl product of the first PCR reaction to barcode the samples for next-generation sequencing. PCR products of the barcoded reaction were verified by running on agarose gel and then extracted using the MinElute Gel Extraction Kit (Qiagen, Cat# 28604) as per manufacturer’s recommendations with a final elution volume of 30 µl in EB buffer. Amplicon libraries were sequenced single-ended (SE) 1x 150 bp on the Illumina NextSeq machine. After demultiplexing, FASTQs were analyzed using CRISPResso2^67^.

### Sequencing data pre-processing

Raw sequencing data for both scATAC-seq and antibody tag libraries were demultiplexed using CellRanger-ATAC mkfastq. For the ATAC data, sequencing reads for all libraries were aligned to the hg38 or hg38/mm10 reference genomes using CellRanger-ATAC count. To eliminate barcode multiplets^68^, all libraries were processed with CellRanger-ATAC v1.2 which utilizes shared Tn5 transposition events to identify and remove barcodes with low tag abundance. Protein tag abundances were estimated using the kallisto, bustools, and kite frameworks^69, 70^. To make the protein tag reads compatible with the kallisto framework processing, we developed an accessory script, ASAP_to_kite.py, that converts fastq files into a format similar to the 10x scRNA-seq format, enabling tag abundance quantification. For CITE-seq data, raw sequencing reads were aligned using CellRanger v3 to the hg38 reference genome. Tag abundances were computed directly using the kallisto, bustools, and kite frameworks^69, 70^.

### Analysis of species mixing experiment

Cells that passed the CellRanger-ATAC knee call were assigned as putative human cells when at least 100 fragments overlapped accessibility peaks in the human reference genome and putative mouse for at least 100 fragments in peaks in the murine reference genome. Similarly, cells were annotated as putative mouse or human cells based on protein abundance based on a minimum count of 100 for human CD29 and 50 for mouse CD29. Doublets were assigned for cells that consisted of less than 95% (ATAC; fragments in peaks) or 90% (protein; CD29 abundance) of the corresponding molecule. All thresholds were determined after evaluation of empirical densities of these measurements. The percent agreement between the multimodal assays was determined using cells that had corresponding labels (mouse, human doublet), which was 97.4% for the pre-SPRI and 97.1% for the post-SPRI experiment. For each experiment, only one cell was observed that was annotated as mouse in one modality and human in the other; the rest of the discrepancies were due to edge cases associated with doublet assignments.

### Complexity analyses

For both protein tag and chromatin complexity estimations, we used the number of unique and duplicate fragments as part of the CellRanger-ATAC (chromatin) and bustools (tag) output as inputs into the Lander-Waterman equation^71^, which estimates the total number of unique molecules present given these two measurements. For chromatin, we used the ’total” and “passed_filters” columns from the singlecell.csv file. For the tag libraries, we converted the corrected bustools file into a tsv file to manually assign and deduplicate reads based on error-corrected barcode, UMI/UBI, and feature assignments. For species mixing experiments, comparisons were performed by selecting the top 1,000 cells ranked by library complexity per condition per species to minimize differences due to variable cell yield (**Extended Data Fig. 1d,e**).

### Resting PBMC analyses

For all analyses in **Fig. 1-3**, gene activity scores, cell clusters, and reduced dimension representations were computed using ArchR^72^ with the default workflow. Visualizations of gene activity scores and protein tag abundances were performed using unsmoothed values after CLR-normalization for the protein tags. From the cell hashing experiment (**Extended Data Fig. 1g**), we assigned putative cell doublet identity using HTODemux^17^ for all barcodes passing the CellRanger-ATAC knee call. Heteroplasmic mtDNA mutations were determined using the mgatk pipeline and variant calling parameters as previously described^14^. The two mutations shown in **Fig. 1f** were selected as they had the highest mean allele frequency among high-confidence heteroplasmic mutations. Violin plots depicting the proportion of mtDNA fragments (**Fig. 2c**) and tag abundances (**Fig. 2d** and **Extended Data Fig. 2c**) were plotted after removing the top 1% of barcodes from the CellRanger-ATAC knee called for each value to minimize the visual impact of artifacts such as cell doublets.

### Bone marrow mononuclear cell analyses

We identified high-quality cells that satisfy three criteria: 1) minimum of a TSS score >4 and 1000 fragments from ArchR^72^, 2) are not doublets based on hashtag oligos / HTODemux^17^, and have less than 10,000 total tags or 50 tags in rat antibodies (cutoffs inferred from density distributions). These steps resulted in 10,928 cells. We then performed LSI, UMAP, and clustering with ArchR using default settings^72^. Annotations of cellular protein tags were performed using CLR-normalized counts among these barcodes. Tag importance was determined after fitting a Random Forest model using the chromatin-derived cluster labels as outcomes and scaled, CLR-normalized protein tag abundances as input features, an approach inspired by the CiteFuse workflow^26^.

Monocytic and erythroid pseudotime was determined using the semi-supervised functionality in ArchR^72^. Protein tag/pseudotime heatmaps were computed by dividing cells into 100 bins, computing means, and then performing a rolling average over 11 consecutive bins as implemented in ArchR^72^. The subset of proteins shown for each lineage were selected such that a) the mean scaled protein tag value exceeded 1 across cells in the trajectory and b) the ratio of means between cells included and excluded in the trajectory exceeded 1. These filtering steps for included proteins were incorporated to minimize the contribution of factors not specifically expressed in these differentiation trajectories. The paired gene score heatmaps were computed using the same procedure but utilizing the single-cell, unsmoothed gene activity scores. Finally, we further restricted the set of genes for the comparison of max protein tag and max gene activity score (**Fig. 4i**) to genes where the protein peaked after 0.25 in the pseudotime directory to eliminate factors associated with multipotent or erythroid-biased progenitors.

Among the cells passing accessible chromatin and protein quality control, 6,797 had a minimum 10x mtDNA coverage, which were considered for downstream mutation analysis. Heteroplasmic mtDNA mutations were determined using the mgatk pipeline and variant calling parameters as previously described^14^. Putative lineage-biased variants were identified using a per-mutation Kruskal Wallis test of association between heteroplasmy and cell lineage, which were assigned to individual cells based on chromatin clusters (see **Supplementary Table 2**).

### Analysis of PBMC stimulation experiments

Control and stimulated CITE-seq cells were filtered using the following criteria: predicted singlets using Scrublet^73^, maximum 10% mitochondrial RNA reads, minimum 500 genes detected, and minimum 1,000 total UMIs observed. Cells were further filtered out if they had excess abundance of total protein tags (>25,000 or 30,000 in control and stimulated conditions, respectively) or tags measured from the rat isotype controls (>55 or 65 in the control and stim, respectively). Similarly, we identified high-quality cells from the ASAP-seq dataset such that each cell had a TSS score exceeding 4 and a minimum of 1,000 fragments. Cells were further filtered out if they had excess abundance of total protein tags (>25,000 in either condition) or tags measured from the rat isotype controls (>75 in either condition). All thresholds for both the ASAP-seq and CITE-seq filtering were determined by evaluating the per-cell empirical density.

We performed two-stage data integration for the ASAP-seq and CITE-seq datasets to preserve the biological effect of the stimulation and residualize differences between the RNA and ATAC assays. First, we created a union of variable genes from the CITE-seq stimulated and control datasets along with genes whose proteins were measured as part of the antibody panel. Using these -2,700 genes, we performed canonical correlation analysis (CCA) between the stimulated ASAP-seq (gene scores) and CITE-seq (RNA abundance) datasets and a second round of CCA between the control ASAP-seq and CITE-seq datasets^74^. For both datasets, we imputed RNA expression for the ASAP-seq objects using transfer anchors as described in Seurat V3^75^. In RNA space for these two merged objects, we performed principal component analysis (PCA), before using Harmony to integrate the stimulated and control integrated datasets^76^. A final dimensionality reduction and clustering using Harmony components was performed to summarize both modalities (ASAP-seq and CITE-seq) and both biological conditions (stimulated and control) in one setting. Finally, the embedding and clustering of ASAP-seq and CITE-seq based on protein tag abundances was performed using the 100 most variable features across the merged ASAP-seq and CITE-seq datasets as inputs to PCA and then Harmony^76^ to account for the technology and stimulation status as two group variables.

In determining the relative changes between chromatin accessibility, RNA, and protein abundance between the stimulated and unstimulated conditions, we generated counts-per-million normalized pseudobulk abundances, which were used to determine the log_2_ fold changes. While these measures were computed for both the B cell and T cell clusters separately, we note that many changes in the B cell population mirrored that of the T cells, which we attributed to low-frequency cell doublets that persisted even after our computational filtering. This inference was based on the presence of markers such as CD4 and CD8 appearing in the B cell clusters, which are markers restricted to T cells and largely unchanged in the stimulation.

Separately, the number of differential peaks, genes, and proteins were computed using a per-peak permutation test^9^, the edgeRQLFDetRate for differential gene expression^77^, and a Mann-Whitney test for the CLR protein abundances. The number of significant differential features (**Fig. 5**) was determined using consistent thresholds of a Benjamini-Hochberg adjusted p-value of 0.01 and a minimum magnitude of log_2_ change exceeding 0.5. The proportion of differential features was computed out of 52,551 peaks, 10,533 genes, and 227 proteins. For accessibility peaks and genes, the universe of those tested were selected based on a mean count per million exceeding 2 across the stimulated and control samples. For the proteins, none of the 71 differentially expressed markers using these criteria were the rat isotype antibodies (known negative controls).

Tag importance (**Extended Data Fig. 5d**) for either the stimulation and cluster identities was determined by fitting two Random Forest models to 1) the cluster labels and 2) the experimental control/stimulation conditions. Again, the scaled, CLR-normalized protein tag abundances were used as input features, an approach inspired by the CiteFuse workflows^26^.

UMAPs showcasing changes in chromatin accessibility, RNA, and protein abundance were consistently displayed using the 2nd and 98th percentiles as minimum and maximum values on the color scale. Cells depicted were displayed in random order. Further, only cells where the modality was directly measured (i.e. chromatin accessibility: ASAP-seq; RNA: CITE-seq; protein: ASAP-seq and CITE-seq) were displayed. Further, no gene smoothing was applied in any display.

### Multiplexed perturbation analyses

From the hashtag count matrix, we assigned perturbation identity using HTODemux^17^. Donor ID per cell was further inferred using popscle, which extends the demuxlet toolkit^78^. High quality cells were determined based on default quality-control criteria using the Signac workflow^75^. Subsequently, these quality controlled cells were used in generating LSI dimensions and the UMAP embedding using ArchR with default settings^72^. Transcription factor accessibility deviation scores were computed using chromVAR with default settings for known human transcription factor motifs, including the inference of the top 100 most variable^47^. Downstream analyses of protein tag abundances were performed on CLR-normalized tag abundances. NFKB-compatible motifs were discovered in chromatin accessibility peaks using the motifmatchr framework as part of the chromVAR^47^ suite of tools.

## ACKNOWLEDGEMENTS

We acknowledge support from the Broad Institute and the Whitehead Institute Flow Cytometry Core facilities. This research was supported by National Institutes of Health grants no. R01 DK103794 (V.G.S.), no. R01 HL146500 (V.G.S.), National Institutes of Health / National Human Genome Research Institute grants no. R21 HG-009748 (P.S.) and no. RM1 HG0110014 (P.S.); Grants-in-Aid by Japan Society for Promotion of Science (JSPS) for Specially Promoted Research no. 16H06295 (S.S.); Japan Agency for Medical Research and Development (AMED) for Leading Advanced Projects for Medical Innovation (S.S.); a gift from the Lodish Family to Boston Children’s Hospital (V.G.S.); the New York Stem Cell Foundation (NYSCF, V.G.S.); the Howard Hughes Medical Institute and Klarman Cell Observatory (A.R.) and the Chan Zuckerberg Initiative / Silicon Valley Community Foundation Human Cell Atlas grant no. HCA3-0000000309 (P.S.); V.G.S. is an NYSCF-Robertson Investigator. C.A.L. is supported by a Stanford Science Fellowship.

## AUTHOR CONTRIBUTIONS

E.P.M. and P.S. conceived and designed the method with input from L.S.L., K.Y.C., and C.A.L. E.P.M., K.Y.C., L.S.L., and P.S. designed experiments with input from C.A.L., V.S. and A.R.. E.P.M., K.Y.C., W.L., P.I.T. and L.S.L. performed experiments. C.A.L. lead data analysis with significant contributions from E.P.M., K.Y.C., and Y.T. A.L.Z.-F., T.-S. H., B.Y., and K.L.N. provided insights and developed key reagents and protocols for protein detection. J.B.W. provided insights and discussions for experimental planning. S.S., L.S.L., V.G.S., A.R., and P.S. each supervised various aspects of the work. E.P.M., C.A.L., K.Y.C., L.S.L., and P.S. drafted the manuscript with input from all other authors.

## COMPETING INTERESTS

P.S. is listed as co-inventor on a patent related to this work (US provisional patent application 62/515-180). C.A.L., L.S.L., V.G.S., and A.R. are listed as co-inventors on a patent related to mtscATAC-seq (US provisional patent application 62/683,502). A.R. is a founder and equity holder of Celsius Therapeutics, an equity holder in Immunitas Therapeutics and until August 31, 2020 was an SAB member of Syros Pharmaceuticals, Neogene Therapeutics, Asimov and ThermoFisher Scientific. From August 1, 2020, A.R. is an employee of Genentech.

## SUPPLEMENTAL TABLES

Supplementary Table 1: Antibodies used in this study Supplementary Table 2: Cell cluster annotations in marrow

**Supplementary Table 3: Dynamic protein tags across monocytic pseudotime Supplementary Table 4: Changes in chromatin, RNA, and cell surface protein in activated T-cells**

Supplementary Table 5: gRNAs and hashtag encodings used for perturbations.

## CODE AVAILABILITY

Custom code to reproduce all analyses and figures is available at https://github.com/caleblareau/asap_reproducibility.

## DATA AVAILABILITY

Data is available at GEO accession GSE156478.

---

**Extended Data Figure 1.**
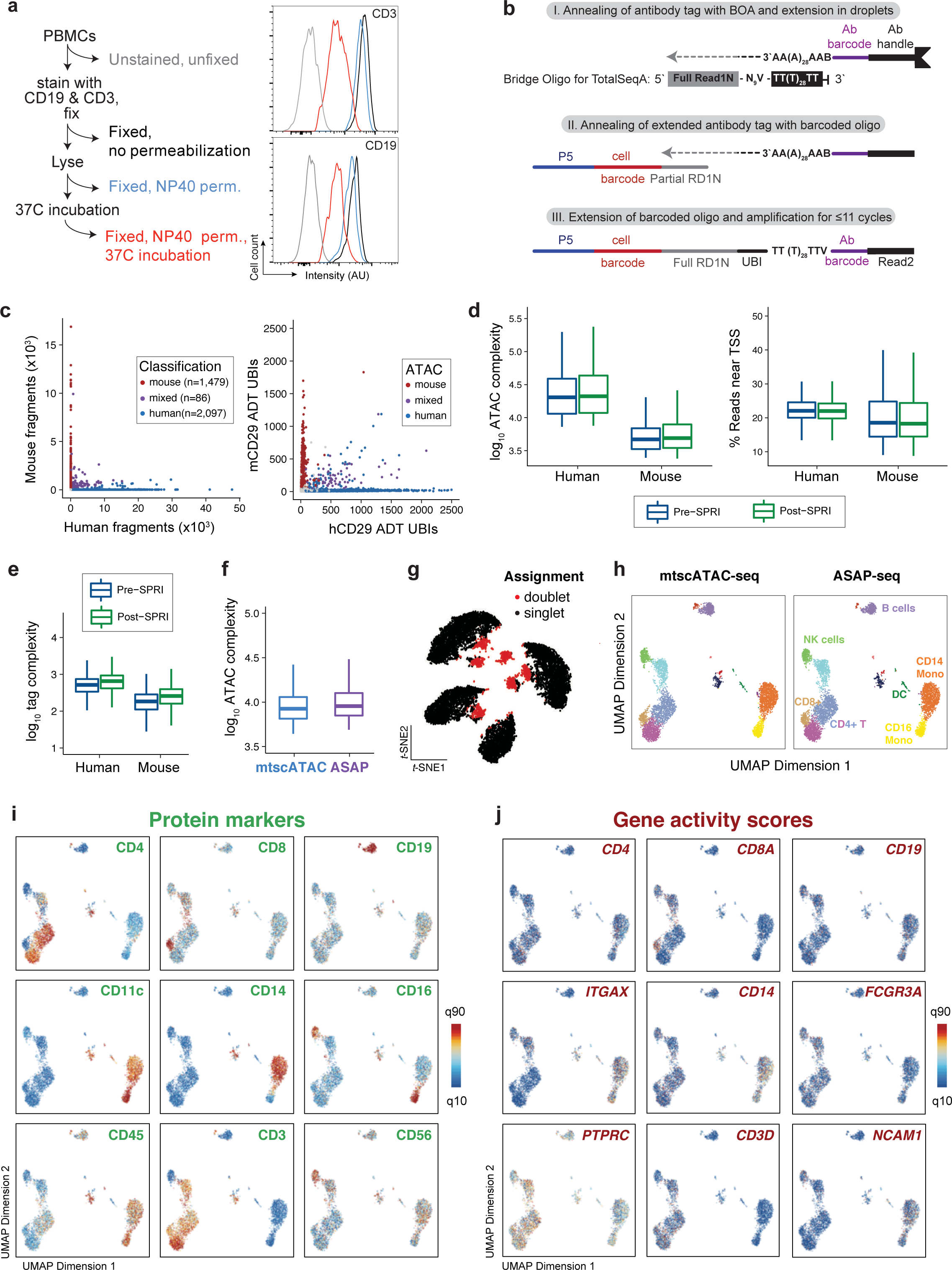
Additional technical and computational validation of ASAP-seq workflows. a. PBMCs were stained with fluorophore-conjugated antibodies and subjected to the ASAP-seq workflow with samples withdrawn at the indicated steps and assessed for fluorophore intensity by flow cytometry. CD3 (top) and CD19 (bottom) signal on fixed cells is hardly affected by permeabilization alone, but after the 37℃ incubation for 1h to mimic the Tn5 transposition reaction, some signal reduction is observed. **b.** Barcoding scheme of TSA tags using the bridge oligo for TotalSeq^TM^-A (BOA). TSA tags do not contain UMIs, so to allow molecule counting, UBIs (N9V) are incorporated via the bridge oligo. **c.** Species mixing experiment as in Figure 1c, using the Post-SPRI approach for tag recovery. Points are colored based on species classification using ATAC fragments. **d.** ATAC library complexity and TSS enrichment for fragments from each species under the two protein-tag library approaches. **e.** Comparison of protein tag complexity between libraries prepared using the pre- and post-SPRI approach. **f.** Comparison of ATAC library complexity between mtscATAC-seq and ASAP-seq. **g.** Two-dimensional embedding of the PBMC hashing data using *t*-SNE. The four major clusters (black) correspond to the four hashing antibodies used to stain the PBMCs. 13,772 cells were recovered and1,396 doublets (red) were detected. **h.** UMAP embedding resolving PBMC cell types based on chromatin accessibility for cells processed by mtscATAC-seq and ASAP-seq. Data for the two different samples were processed together using cell ranger-atac aggr before dimensionality reduction. **i.** Selected protein markers (left) and corresponding gene score activities (right) superimposed on the ATAC-clustered PBMCs (for the ASAP-seq sample) as in (**h**).

**Extended Data Figure 2.**
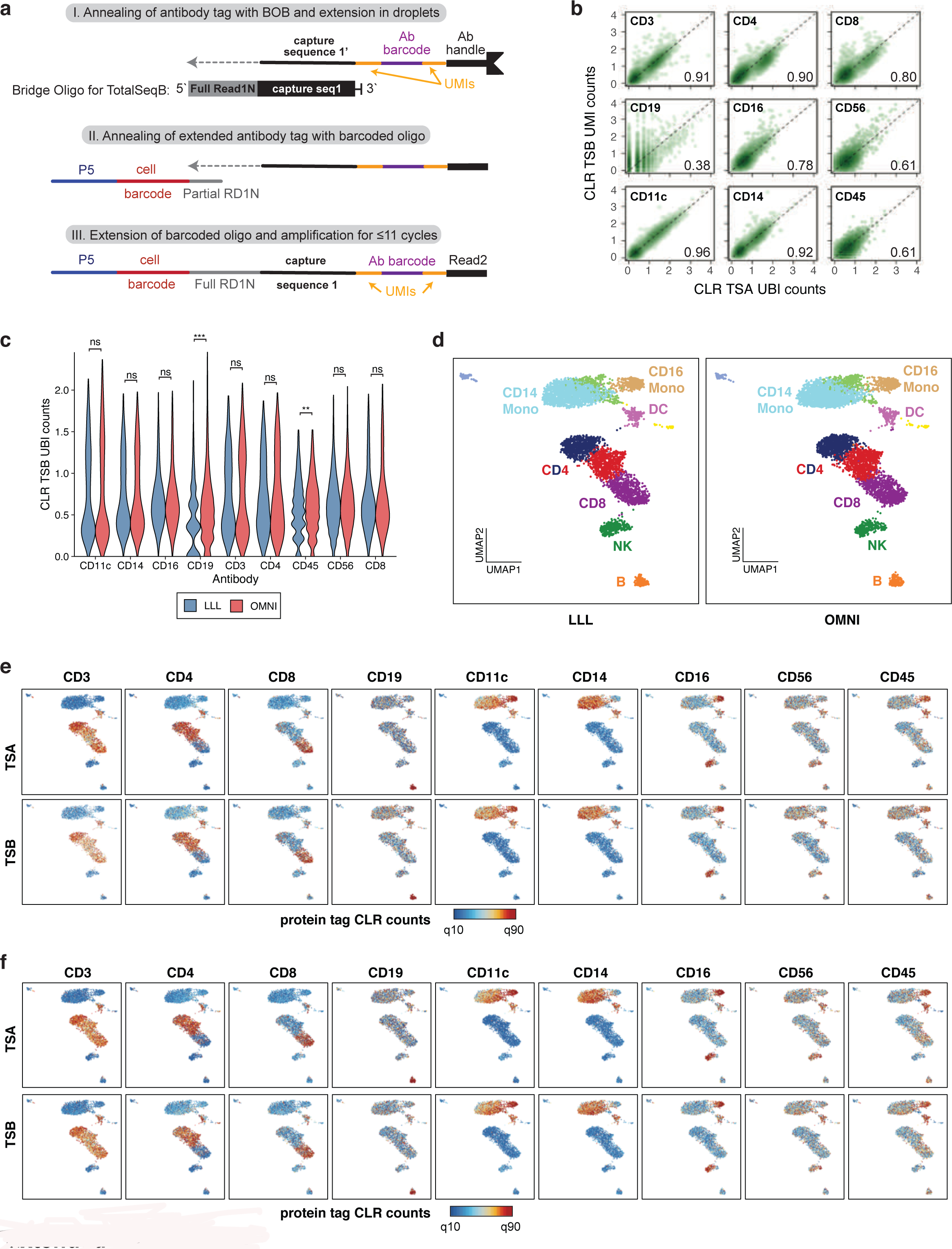
Additional validation and comparison of modular ASAP-seq workflows. a. Barcoding scheme of TSB tags using the bridge oligo for TotalSeqB (BOB). TSB tags contain UMIs (encompassing the antibody barcode), negating the requirement for a UBI on the bridge oligo. **b.** Pairwise comparison of centered log-ratio (CLR) normalized TSA and TSB counts under OMNI lysis conditions (n=5,236 cells). Counts were collapsed for unique molecules using UBIs (TSA panel) or UMIs (TSB panel). **c.** Comparison of CLR normalised TSB counts under the two lysis conditions. Statistical comparisons are Wilcoxon rank sum test with Bonferroni adjusted p-values (ns = not significant; \**p_adj_* < 0.05; ** *p_adj_* < 0.01; *** *p_adj_* < 0.001). **d.** UMAP embedding and cluster annotation of the LLL (n=5,236) and OMNI (n=4,748) processed cells. Data for the two different samples were processed together using cell ranger-atac aggr before dimensionality reduction. **e.** TSA and TSB CLR counts projected on the LLL embeddings. **f.** TSA and TSB CLR counts projected on the OMNI embeddings.

**Extended Data Figure 3.**
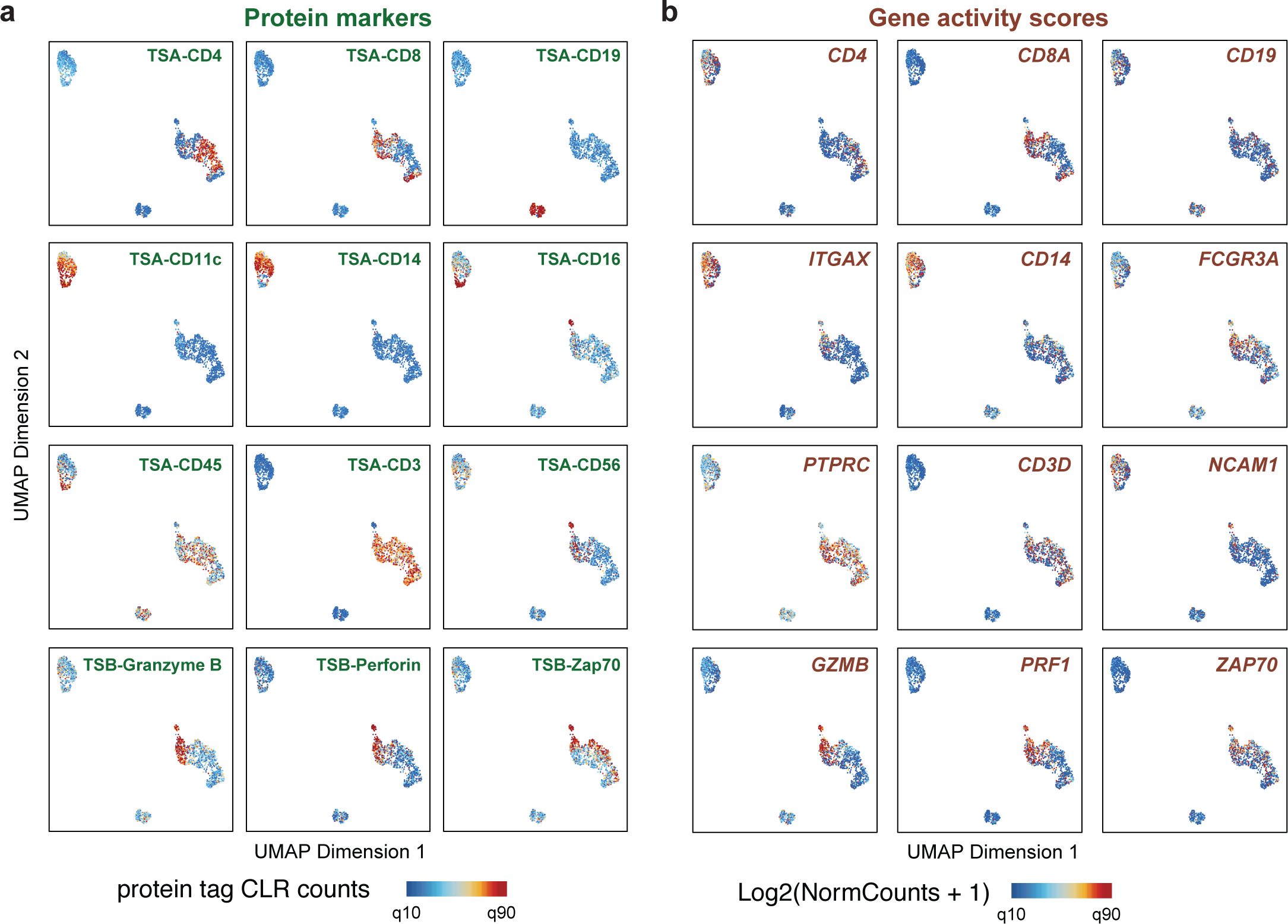
Supporting information for intracellular ASAP-seq workflow. a,b. Selected protein markers (**a**) and corresponding gene activity scores (**b**) superimposed on the ATAC-clustered PBMCs from the intracellular staining experiment (see Figure 3a).

**Extended Data Figure 4.**
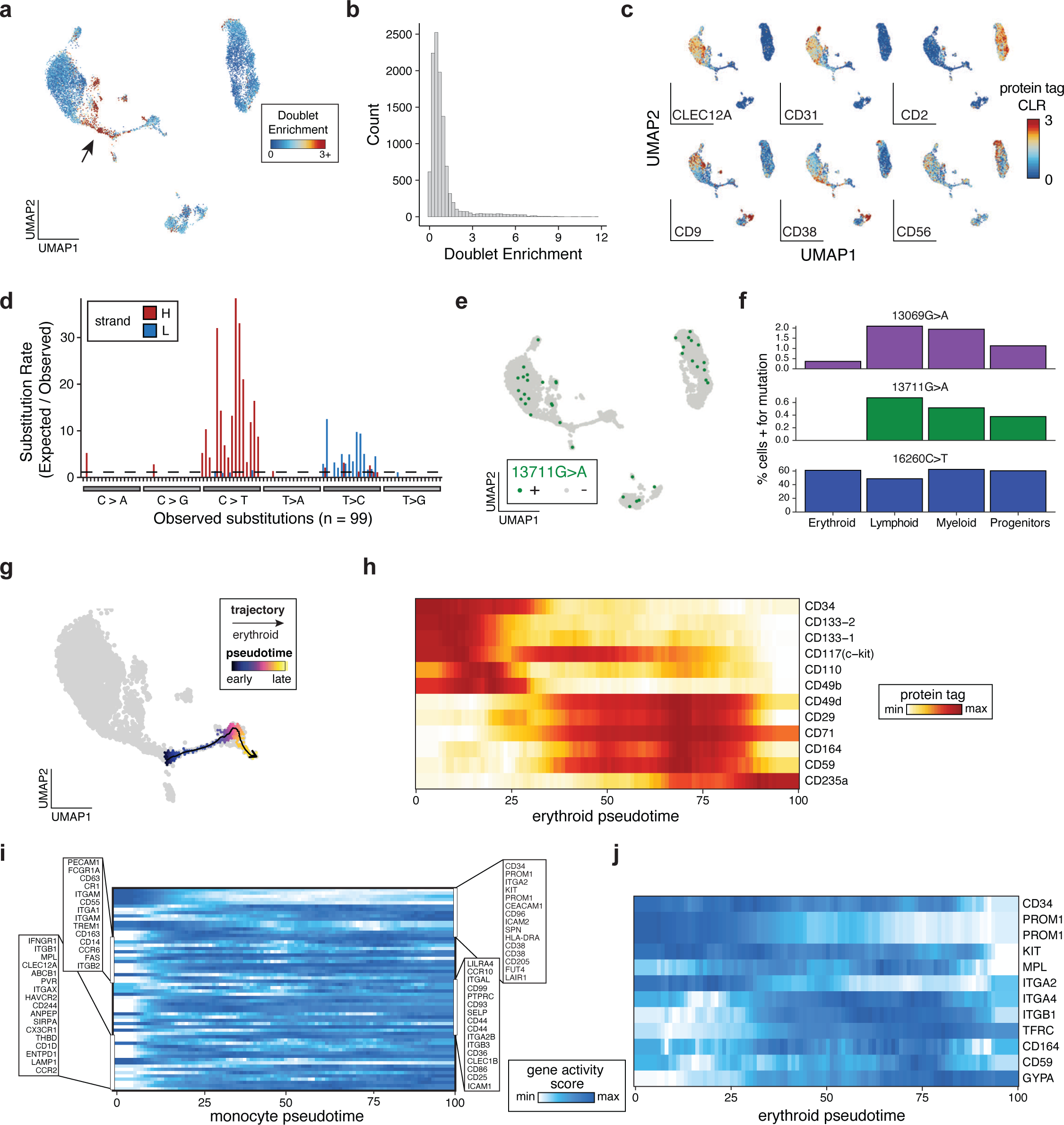
Supporting information for ASAP-seq bone marrow analyses. a. Annotation of reduced dimension space with the Doublet Enrichment score from ArchR. Arrow indicates the monocytic progenitor population. **b.** Histogram of scores from panel (**a**). **c.** Feature plots for six additional antibody tags in the reduced dimension space. **d.** Substitution rate (observed over expected) of mgatk-identified heteroplasmic mutations (y axis) in each class of mononucleotide and trinucleotide change resolved by the heavy (H) and light (L) strands of the mitochondrial genome. **e.** Projection of 13711G>A in single cells; threshold for + was 5%. **f.** Distribution of observed mtDNA mutations in cells among major cell lineages. **g.** Expression of chromatin activity scores along the monocytic developmental trajectory for genes encoding proteins shown in Figure 4h. **h.** Developmental trajectory of erythroid differentiation using semi-supervised pseudotime analysis. **i.** Expression of select cell surface markers along the erythroid developmental trajectory highlighted in (**h**). Rows are min-max normalized. **j.** Expression of chromatin activity scores along the erythroid developmental trajectory for genes encoding proteins shown in (**i**).

**Extended Data Figure 5.**
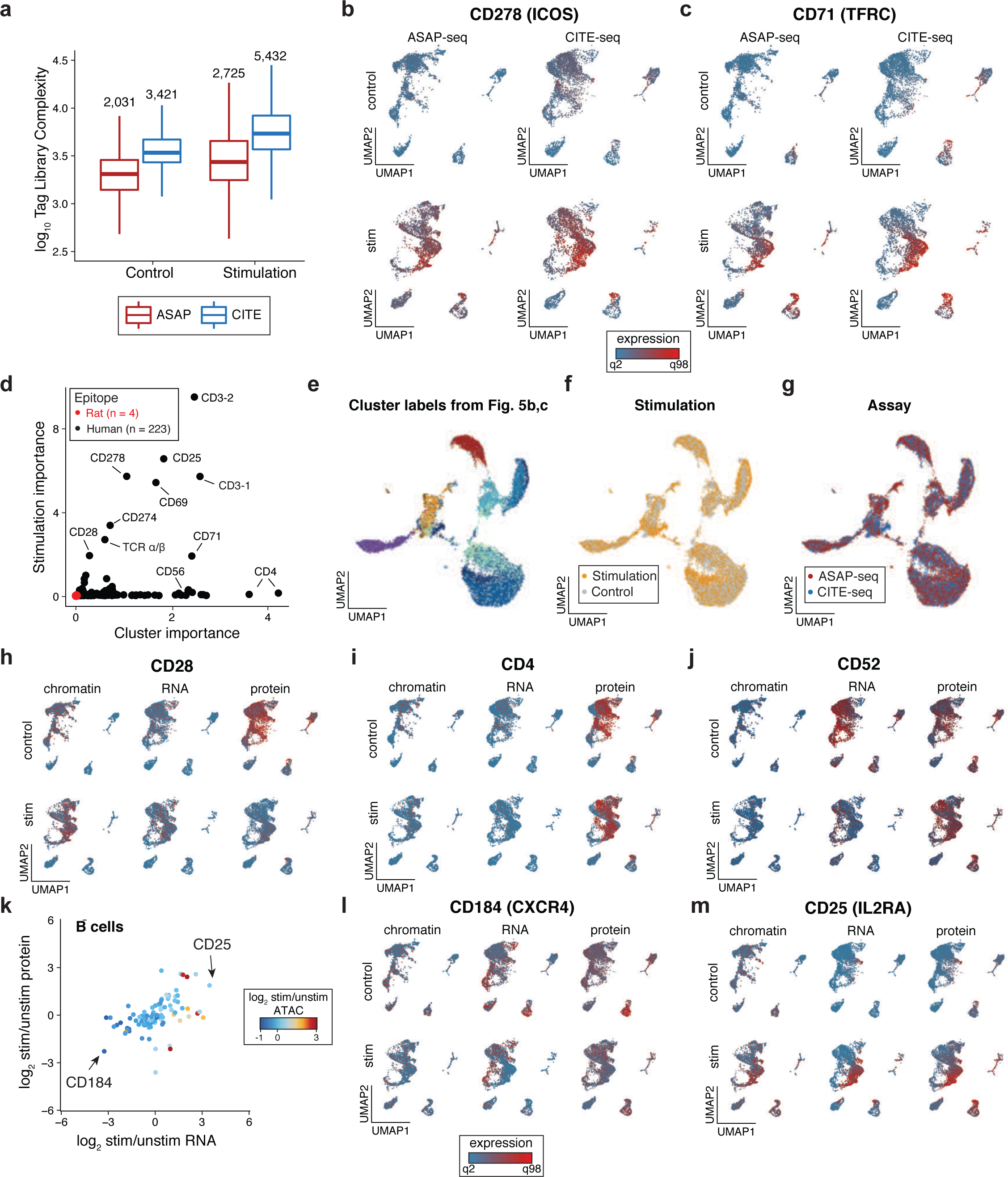
Supporting information for combined ASAP-seq and CITE-seq readouts. a. Antibody tag complexity per condition and technology. Median tag complexity is 1.7-2x higher in CITE-seq compared to ASAP-seq and 1.3-1.6x higher in stimulation compared to control sample. **b,c.** Cellular distribution of protein tags measured by ASAP-seq (left) and CITE-seq (right) for control (top) and stimulated conditions (bottom) for, (**b**) CD278 (*ICOS*) and (**c**) CD71 (*TFRC*). **d.** Protein tag measurement importance in predicting cell cluster and stimulation from two different Random Forest models. Negative controls (rat epitopes) are shown in red. **e-g.** ASAP-seq and CITE-seq data co-embedding utilizing protein abundances. Cells are highlighted by (**e**) chromatin/RNA cluster identity, (**f**) stimulation condition and (**g**) technology assayed. **h-j.** UMAPs of chromatin accessibility, mRNA expression, and surface protein levels for (**h**) CD28, (**i**) CD4, and (**j**) CD52. **k.** Summary of changes in chromatin accessibility, gene expression and surface protein abundance for 103 expressed genes in B cells following T cell stimulation. **l,m.** UMAPs of chromatin accessibility, mRNA expression, and surface protein levels for genes with differential expression in B cells, including (**l**) CD184 (*CXCR4*) and (**m**) CD25 (*IL2RA*).

**Extended Data Figure 6.**
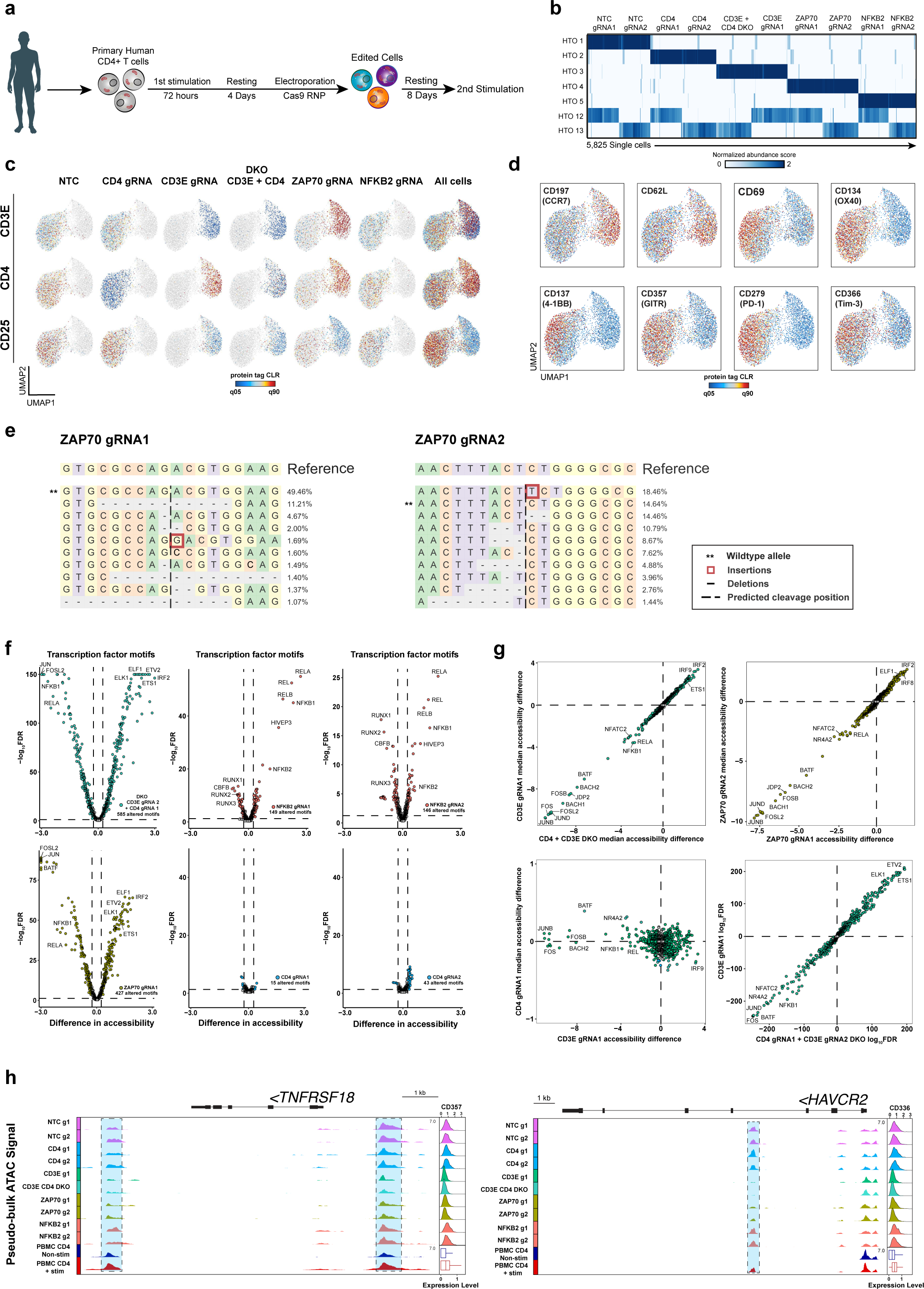
Supporting information for ASAP-seq based decoding of perturbations in primary T cells. a. Schematic for CRISPR perturbation experiment in primary human T cells. CD4+ T cells from healthy donors were isolated and stimulated for 72 hours, followed by a resting period of four days to enable expansion. On Day 7, cells were electroporated with Cas9 RNPs and then rested for an additional 8 days before secondary stimulation. **b.** Heatmap of cell demultiplexing with hashing antibodies, indicating normalized abundance of each hashtag. **c.** Assessment of the effect of CRISPR perturbations on three indicated protein surface markers. **d.** UMAP embedding overlaid with expression of the eight indicated surface protein markers. **e.** Allele-specific CRISPR editing outcomes for ZAP70 gRNA1 (left) and ZAP70 gRNA2 (right). The wildtype allele is indicated by **. **f.** Volcano plots showing transcription factor motifs with significantly changed chromatin accessibility profiles between NTC cells and the indicated gRNAs (FDR <= 0.05, chromVAR accessibility change >= 0.25). **g.** Correlation of chromVAR median accessibility changes or FDR (bottom right panel) between the indicated gRNAs. **h.** Genomic tracks of *TNFRSF18* and *HAVCR2* loci with corresponding CLR-normalized protein abundance ridge plots. CLR-normalized protein abundance from the PBMC stimulation experiment is indicated by the corresponding boxplots. Differentially accessible regions are highlighted in blue.

## Notes

https://github.com/caleblareau/asap_reproducibility

### Competing Interest Statement

COMPETING INTERESTS
PS is listed as co-inventor on a patent related to this work (US provisional patent application 62/515-180). CAL, LSL, VGS, and AR are listed as co-inventors on a patent related to mtscATAC-seq (US provisional patent application 62/683,502). A.R. is a founder and equity holder of Celsius Therapeutics, an equity holder in Immunitas Therapeutics and until August 31, 2020 was an SAB member of Syros Pharmaceuticals, Neogene Therapeutics, Asimov and ThermoFisher Scientific. From August 1, 2020, A.R. is an employee of Genentech.

